# Cancer associated talin point mutations disorganise cell adhesion and migration

**DOI:** 10.1101/2020.03.25.008193

**Authors:** Latifeh Azizi, Alana R. Cowell, Vasyl V. Mykuliak, Benjamin T. Goult, Paula Turkki, Vesa P. Hytönen

## Abstract

Talin–1 is a key component of the multiprotein adhesion complexes which mediate cell migration, adhesion and integrin signalling and has been linked to cancer in several studies. We analysed talin–1 mutations reported in the COSMIC (Catalogue of Somatic Mutations in Cancer) database and developed a bioinformatics pipeline to predict the severity of each mutation. These predictions were then assessed using biochemistry and cell biology experiments. With this approach we were able to identify several talin–1 mutations affecting integrin activity, actin recruitment and Deleted in Liver Cancer 1 localization. We explored potential changes in talin–1 signalling responses by assessing impact on migration, invasion and proliferation. Altogether, this study describes a pipeline approach of experiments for crude characterization of talin–1 mutants in order to evaluate their functional effects and potential pathogenicity. Our findings suggest that cancer related point mutations in talin–1 can affect cell behaviour and so may contribute to cancer progression.

## Introduction

For cells to maintain homeostasis and co–operate within tissues, they need to dynamically interact with the extracellular matrix (ECM). In recent years the role of the microenvironment has become increasingly recognised^1^ and disturbances between cell–ECM interactions, intracellular signalling events, and signals derived from the ECM have been shown to contribute to cancer progression. Talin is a major component of focal adhesions (FAs), responsible for mediating the link between the ECM via integrins and the actin cytoskeleton (Fig. 1A). Talin is a large ~250 kDa mechanosensitive protein consisting of an N–terminal FERM head domain (F0, F1, F2, F3; residues 1–405) followed by a linker (~80aa) and ~2000aa rod region comprised of 13 domains (R1 to R13) ending in a C–terminal dimerisation domain (DD)^2^ (Fig.1B). The FERM domain interacts with the membrane–proximal NPxY motif of beta integrin tail and the negatively charged plasma membrane^3, 4^ The rod domain contains two F–actin binding sites (ABS2 and ABS3)^5–7^, 11 vinculin binding sites (VBS)^8^ and binding sites for regulatory proteins such as RIAM, KANK^9, 10^ and the tumour suppressor DLC–1^11, 12^ (Fig. 1B). Studies have shown that ABS3 is essential for FA assembly^13^, whereas ABS1 and ABS2 have a reinforcing role^14, 15^.

**Figure 1:**
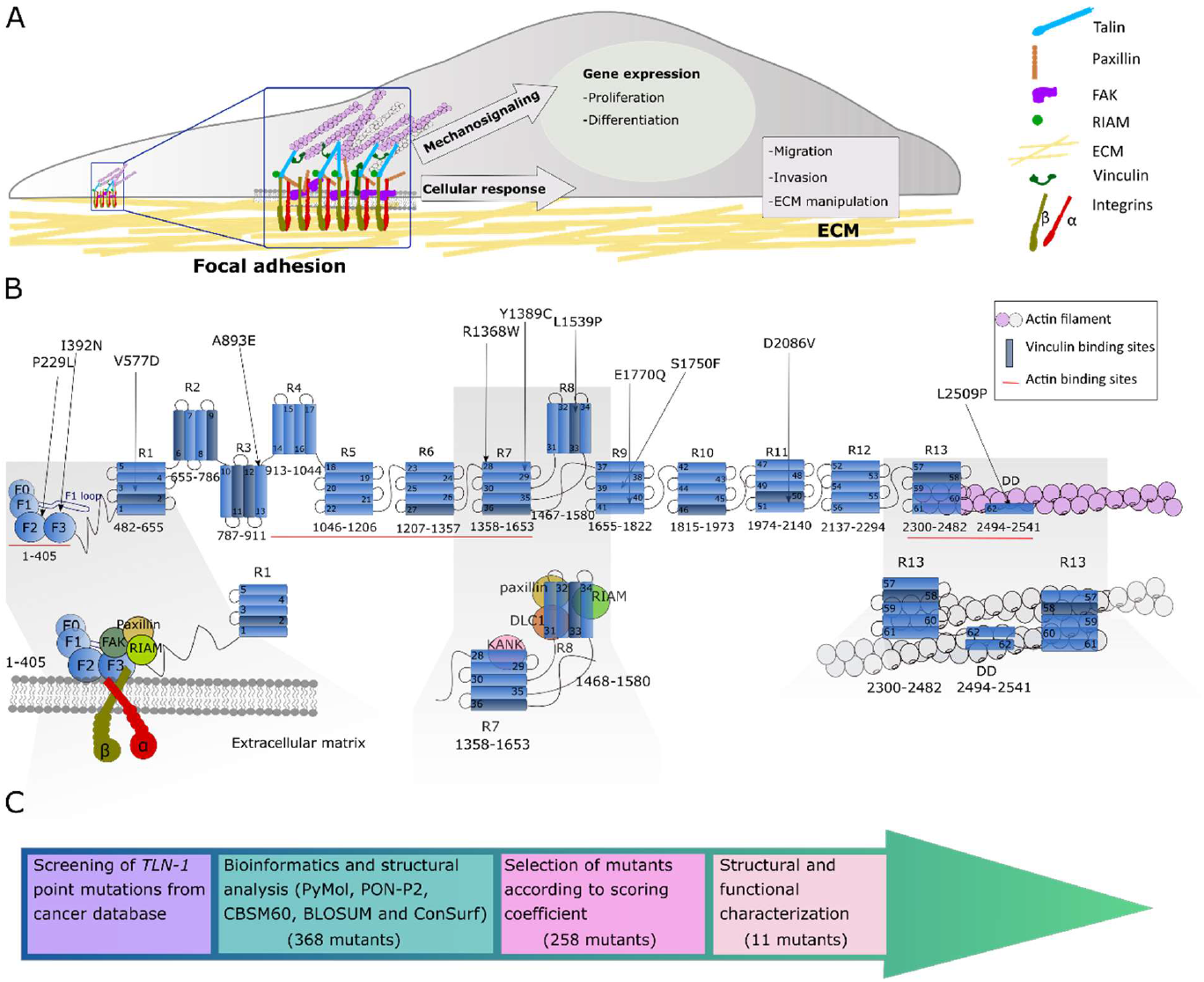
Analysis of talin–1 mutations found in the COSMIC database. A) Schematic representation of focal adhesion role on cell function. B) Schematic representation of talin–1 and the positions of the selected missense mutations. Bottom: cartoon of the talin head complexes that form with integrin, FAK, paxillin and RIAM (left), talin R7–R8 complexes with KANK, DLC–1, RIAM and paxillin (middle) and the R13–DD actin cytoskeletal linkage (right). C) Flowchart showing the study pipeline to investigate TLN–1 mutations from COSMIC database.

Talin is the main scaffold protein in focal adhesions which form at the leading edge of a polarised cell. Talin links the intracellular tails of integrins to the actin cytoskeleton and mechanical forces exerted on talin can disrupt and reveal binding sites leading to formation of mature multiprotein FA complexes^16–18^. These are dynamic processes regulated by a complex signalling network, gathering information from intracellular and extracellular events. The mechanical properties of the ECM are reflected by intracellular changes via the FAs and actomyosin network, having a direct effect on cell behaviour, such as cell shape, migration and proliferation^19, 20^ (Fig.1A).

Talin–1 overexpression has been shown to correlate with increased invasion and decreased survival with oral squamous cell carcinoma^21^ as well as migration, invasion and anoikis resistance in prostate cancer cells^22^. Loss of talin–1 leads to diminished *in vivo* metastasis of prostate cancer cells via FAK–Src complexes and AKT kinase signalling^22^ Talin–1 knockdown was also shown to significantly reduce the proliferation, migration and invasion of colorectal cancer cell line. This was associated with downregulation of several factors which are involved in the epithelial–to–mesenchymal transition, such as STAT3 and vimentin^23^. Conversely, downregulation of talin–1 has also been shown to promote hepatocellular carcinoma progression^24^.

The COSMIC (Catalogue Of Somatic Mutations In Cancer) database^25^ contains exon sequencing data of human cancers and provides a vast resource of somatic mutation information. In light of talins integral role in regulating cell behaviour, integrin adhesion signalling and its connection to cancer progression, we sought to explore how cancer–associated talin mutations may alter talin function and the behaviour of cells.

Here we provide a pipeline approach in order to evaluate the effect of talin–1 mutations. First, we applied a set of bioinformatic prediction tools in order to rank each mutation based on its location and predicted effects on protein structure. We used the scoring function to select mutations with the potential to disrupt talin function and/or affect its interactions with known binding partners for further characterisation. Next, we assessed the mutations effects via computational methods, and biochemical and cell biological analysis was performed to determine the mutations effects on known interactions. Lastly, we studied the effects of mutations on essential cellular behaviours, such as cell migration, invasion and proliferation. This pipeline would be amenable to other proteins and so provides a way to convert the COSMIC information into biological insights.

## Results

### Screening of *TLN–1* mutations from COSMIC database

To investigate functional consequences of talin–1 point mutations, 368 talin–1 mutations in COSMIC database (accessed January 2017) were evaluated, and 258 missense mutations were further screened using bioinformatic tools (Fig.1C). We investigated each of the 258 point mutations individually and determined the position of the mutation within the talin structure (Fig.1B; Fig.S1). The pathogenicity of the amino acid substitutions were predicted using the PON–P2–algorithm^26^. We used a BLOSUM62 substitution matrix^27^ and CBSM60 matrix^28^ to obtain a numeric penalty for each amino acid deviation and to predict the protein structure/function, respectively. Next we investigated if the mutations cause changes in amino acid polarity as the stability of talin domains are strongly dependent on hydrophobic effect^29, 30^. The degree of evolutionary conservation of the amino acids in the talin sequence was investigated using ConSurf^31^. Finally, we evaluated if the mutation is close to known ligand–binding sites. All these factors were used to build a scoring coefficient and the weight for each variable was obtained using the manual iteration process described in the Supplementary material.

Based on this analysis, eleven mutations were selected for further investigation with the E1770Q mutation also included despite a lower score due to its location in the previously defined talin autoinhibition site^32, 33^ (Table 1;Fig.1B).

**Table 1.** The list of talin–1 point mutations selected from the COSMIC database. In each column a normalised value close to 1 predicts defects in protein function. BH=mutation located between helices and DD=dimerisation domain. CBSM6O=conformation-specific amino acid substitution matrix. BLOSUM62=BLOcks SUbstitution Matrix. The recurrence value has been updated September 13^th^, 2020.

| Talin-1 mutant | Recurrence | Primary tissue | Histology | Domain | Location | PON-P2 probability of pathogenicity | Normalised BLOSUM 62-matrix score | Normalised CBM60 matrix score | Normalised ConSurf score | Final score |
| --- | --- | --- | --- | --- | --- | --- | --- | --- | --- | --- |
| P229L | 2 | Skin | Carcinoma | F2 | BH/buried | 0.94 | 1 | 1 | 1 | 8.20 |
| I392N | 1 | Pancreas | Carcinoma | F3 | Buried | 0.86 | 1 | 0.83 | 1 | 8.95 |
| V577D | 1 | Liver | Carcinoma | R1 | Buried | 0.94 | 1 | 1 | 0.60 | 8.39 |
| A893E | 3 | Central nervous system/pituitary | Glioma/Cranio pharyngioma | R3 | BH/buried | 0.95 | 0.75 | 0.33 | 0.60 | 7.63 |
| R1368W | 2 | Hematopoietic and lymphoid tissue /large intestine | Lymphoid neoplasm /carcinoma | R7 | Surface | 0.98 | 1 | 1 | 1 | 7.75 |
| Y1389C | 2 | Liver | Carcinoma | R7 | Buried | 0.90 | 0.87 | 0.67 | 1 | 8.58 |
| L1539P | 1 | Liver | Carcinoma | R8 | Buried | 0.98 | 1 | 1 | 0.60 | 8.45 |
| S1750F | 1 | Skin | Malignant melanoma | R9 | Buried | 0.89 | 0.87 | 0.5 | 0.60 | 7.55 |
| E1770Q | 1 | Breast | Carcinoma | R9 | Surface | 0.81 | 0.37 | -0.33 | 1 | 6.01 |
| D2086V | 1 | Breast | Carcinoma | R11 | Surface | 0.96 | 1 | 1 | 1 | 7.71 |
| L2509P | 1 | Large intestine | Carcinoma | DD | DD /Surface | 0.96 | 1 | 1 | 0.64 | 7.46 |

We next evaluated the selected COSMIC mutations against the 1000 Genomes Project database, which is a large database of human genetic variant data^34^. The R1368W mutation from our selection was the only one additionally found in the 1000 Genomes database (accessed September 8^th^, 2020), indicating that this mutation has also been found in apparently healthy individuals.

### Molecular Dynamics simulations suggest that the majority of mutations destabilise the talin–1 subdomains

To investigate how mutations affect talin–1 subdomain stability, we employed Molecular Dynamics (MD) simulations. Non–equilibrium alchemical MD simulations were conducted to predict the free energy changes in subdomain stability upon mutation (Fig.S2A). All mutations were analysed except those that involve proline (P229L, L1539P and L2509P), which is not supported for the analysis. The majority of the mutations showed destabilisation of the corresponding subdomains, where the strongest destabilisation effect was for mutations V577D in R1 (52.15±13.71 kJ/mol) and I392N in F3 (38.28±1.27 kJ/mol). Strong destabilisation was also showed for Y1389C in R7 (20.60±4.09 kJ/mol). Weaker, but still significant destabilisation was observed for mutations D2086V in R11 (12.96±0.82 kJ/mol), R1368W in R7 (12.63±0.27 kJ/mol), and A893E in R3 (10.5±1.91 kJ/mol). Mutation E1770Q in R9 (−4.17±0.35 kJ/mol) showed mild stabilisation effect, and S1750F also in R9 (2.28±2.33 kJ/mol) did not cause any significant change in the subdomain’s stability.

For more detailed characterisation, we next selected mutations closest to sites with known interaction partners and employed equilibrium MD simulations. In particular, we selected: P229L and I392N due to their proximity to the integrin binding site in F3; R1368W, Y1389C, and L1539P within the R7–R8 fragment which is known to interact with paxillin, RIAM, KANK and DLC–1; and L2509P which is situated within the DD and the C–terminal actin binding site, ABS3^7^ (Fig. 1C).

The P229L mutant is located at the interface between F2–F3. Analysis of F2–F3 inter-subdomain binding energy for F2–F3 WT and P229L mutant suggested that the mutation did not alter the F2–F3 interactions significantly (Fig.S2B). Mutation of Isoleucine 392, which is located inside the hydrophobic core of the F3 subdomain, to Asparagine (I392N) leads to replacement of the hydrophobic isoleucine side chain with a hydrophilic side chain. This strongly destabilises the F3, and equilibrium MD suggests that water molecules can penetrate the destabilised F3 fold (Fig.2A).

**Figure 2:**
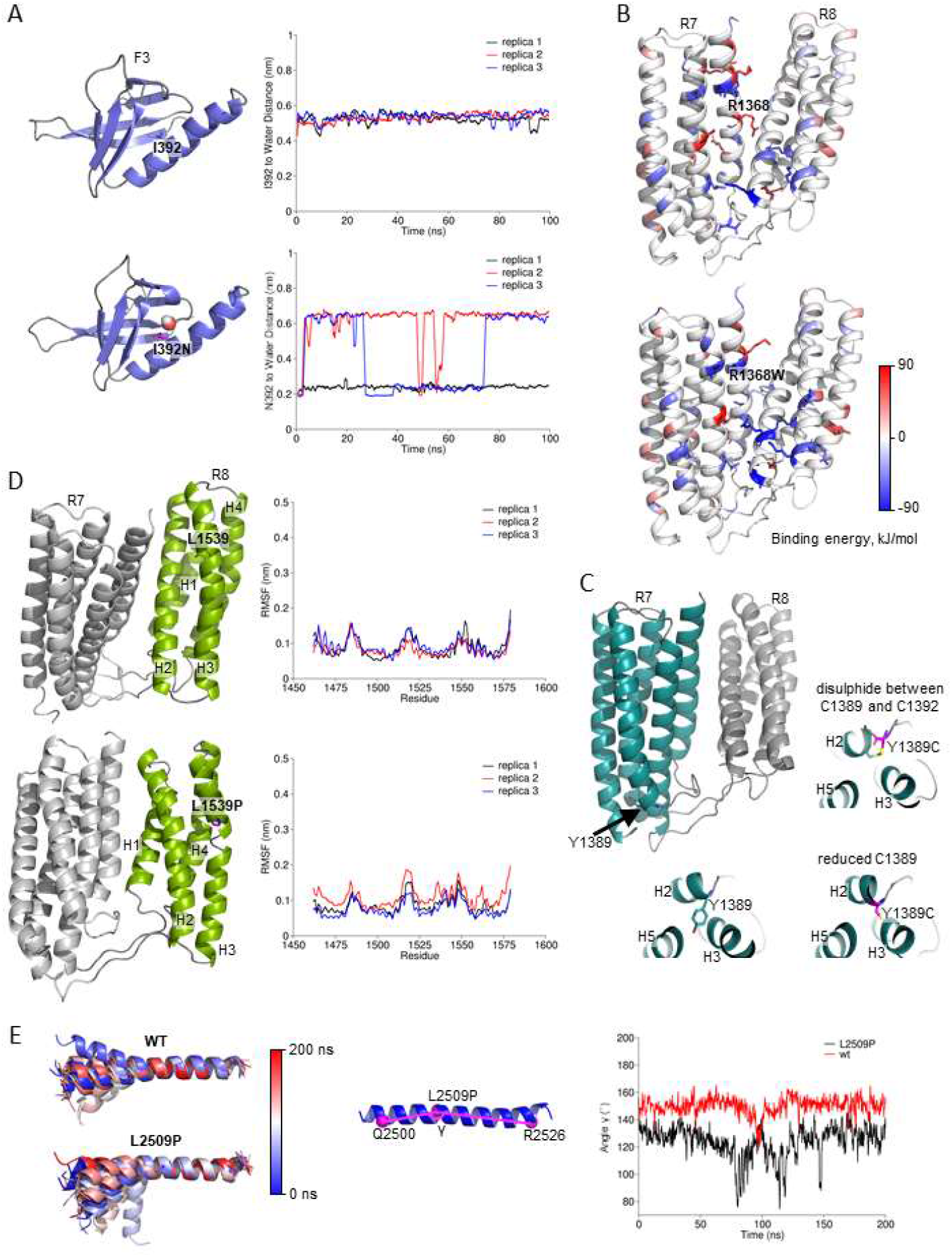
MD simulation analysis of the effects of cancer–associated mutations on talin domains. A) Structure snapshots of the F3 subdomain for WT and I392N mutant. Residue 392 and a water molecule inside the F3 domain in the I392N mutant simulation are shown. The plots show the distance between the residue 392 side chain and the closest water molecule as function of the MD simulation time in WT (top) and I392N mutant (bottom). Three 100 ns replicas are shown. The distance of ~0.6 nm in the WT indicates that the closest water molecule is located at the F3 surface, while in the I392N mutant the distance ~0.2 nm indicates that the nearest water molecule has penetrated the F3 fold. B) R7–R8 inter–domain binding energy distribution for WT and R1368W predicted using MM-PBSA. Residue R1368W and other that have contribution to the inter–domain binding energy lower than −50 kJ/mol and higher than 50 kJ/mol are shown as sticks. C) Visualisation of Y1389C mutation in R7–R8 structure. D) R7–R8 structure snapshots captured at 100 ns of MD for WT and L1539P (in R8); the R7–R8 was used in the simulations, but the Root Mean Square Fluctuation (RMSF) analysis was performed for R8 only. Proline breaks the secondary structure and increase the flexibility of the domain, which reflects RMSF. R7 is shown in grey and R8 in green. E) MD simulations for DD helix showing increased flexibility of the helix caused by L2509P compared to the WT. Superposition was performed using C–alpha atoms of residues 2510 to 2529. The angle (γ) in L2509P mutant and WT in a single helix measured as a function of time.

MD simulations carried out for R7–R8 WT and associated mutants (R1368W, Y1389C, and L1539P) gave the unexpected finding that the R7 and R8 subdomains interact over the course of 100 ns MD (Fig.S2C). Whilst this interaction was detected for all four variants, the analysis of the R7–R8 interdomain binding energy for WT and R1368W mutant suggested that the R1368W mutation enhances this transient interaction between R7 and R8 (Fig.2B, Fig.S2B). The Y1389C mutation is positioned close to a cysteine residue C1392, raising the possibility of a potential intradomain disulphide bond forming. To explore this, two forms of the Y1389C mutation were analysed in equilibrium MD, one where C1389 forms a disulphide bond with C1392, and one where it remains reduced (Fig.2C). Although the mutation strongly destabilises R7 by 20.60±4.09 kJ/mol, equilibrium simulations did not show significant structural changes in R7 in either of these scenarios. MD simulations revealed that the L1539P mutation effectively breaks the structure of helix three (H3) in the R8 subdomain (Fig.2D), and secondary structure analysis indicated clear disruption of H3 in the L1539P–mutated R8 when compared to WT (Fig.S2D). Finally, the L2509P mutation in the DD helix breaks its helical structure and causes higher flexibility of the DD (Fig.2E).

### Biochemical characterisation indicates that Y1389C decreases R7–R8 domain stability and L2509P leads to loss of dimerisation

We selected mutations located in the previously identified binding sites for known talin interactors and which showed strong destabilization of the domain in free energy calculations (I392N, R1368W, Y1389C, L1539P and L2509P) for further biochemical characterisation. For this purpose, we recombinantly expressed talin head fragment 1–405 with mutation I392N, the R7–R8 fragment 1355–1652 versions containing mutations R1368W, Y1389C, L1539P and the R13–DD fragment 2300–2541 with mutation L2509P in *E. coli*. In addition, we expressed the corresponding wildtype protein fragments. We found that the talin head fragment containing I392N mutation purified as two separate fragments and the fragment sizes suggested that cleavage had occurred within the vicinity of the mutated site. A similar cleavage was observed previously with a recombinantly expressed, G340E (equivalent to G331E in mouse talin–1) mutant in fly talin F3 domain^35^ suggesting that destabilisation of F3 leads to exposure of proteolytically sensitive regions that are degraded by *E. coli* proteases. Adding a cocktail of proteolysis inhibitors during the lysis and purification did not help sufficiently to enable production of intact mutant for biochemical analysis. Additionally, R7–R8 containing the mutation L1539P in R8 expressed poorly with evidence of heavy aggregation, making it impossible to obtain protein concentrations high enough for further biochemical analyses. Therefore, we did not pursue further with the biochemical characterization of mutants I392N and L1539P. Wildtype R7–R8, R7–R8 R1368W and R7–R8 Y1389C expressed well and had clear monomeric peaks on size exclusion chromatography (SEC) (Fig.3A). We used Nuclear Magnetic Resonance (NMR) and Circular Dichroism (CD) to provide biophysical insight into the structural effects of these mutations. Comparison of the ^15^N–HSQC spectra of the R7–R8 WT and R7–R8 R1368W mutant revealed similar peak dispersion in both spectra, with only minor shifts in peak positions that occur as a result of the altered sidechain suggesting that R7–R8 R1368W does not significantly affect talin R7–R8 structure (Fig.S4A). In support of this, CD analysis of the R7–R8 WT and R7–R8 R1368W proteins yielded similar spectra confirming similar secondary structure, and no change in thermal stability (Fig.3B,C).

**Figure 3.**
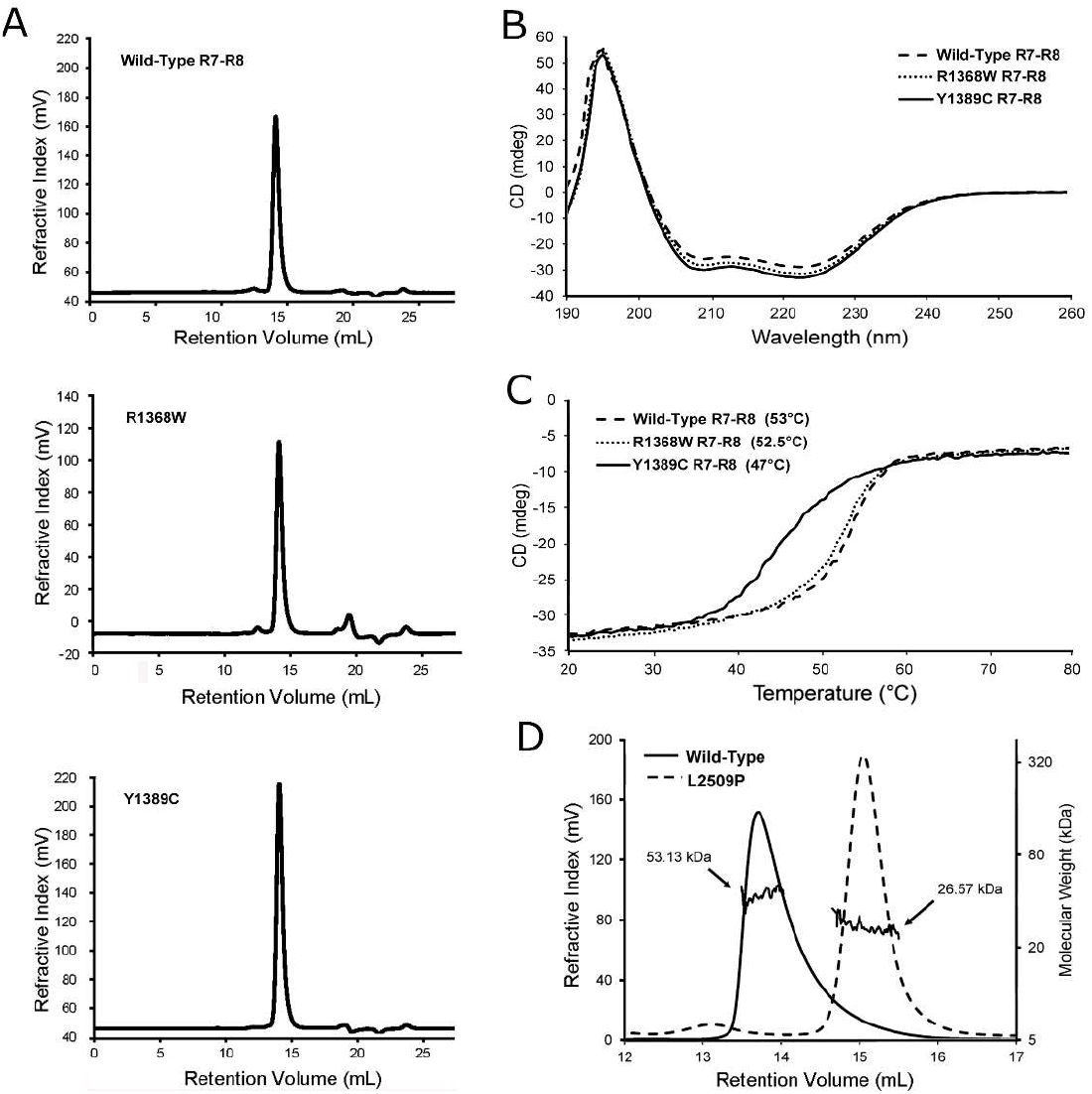
Influence of point mutations on biophysical properties of talin. A) Full SEC profile of R7–R8 WT, R7–R8 R1368W and R7–R8 Y1389C showing the monomeric state. B–C) CD analysis of R7–R8 WT, R7–R8 R1368W and R7– R8 Y1389C. (B) CD spectra of each mutant. (C) Melting temperature curves; the melting temperature of each protein is shown. D) SEC–MALS analysis of R13–DD WT and R13–DD L2509P showing that the R13–DD L2509P is monomeric. The molecular weight obtained from MALS is shown for each peak.

In contrast, the R7–R8 Y1389C mutation caused a striking 6.5°C reduction in the thermostability of R7–R8 (45.5°C compared with 52°C for the wildtype) assessed using CD (Fig.3C). Mutation to cysteine introduces the possibility of a potential to form a disulphide bond with the adjacent cysteine C1392. To test whether the thermal destabilization was due to intramolecular disulphide bond, we performed this analysis in the presence and absence of dithiothreitol and saw no difference between the melting point (data not shown). Unfortunately, NMR analysis of this mutant was not possible, potentially linked to reduced solubility of the mutated protein making concentrating the sample difficult.

Finally, L2509P, located in the talin dimerisation domain, was predicted to disrupt talins ability to dimerise and thus alter actin binding as shown in previous research^7^. Size exclusion chromatography with Multi–Angle Light Scattering (SEC–MALS) showed that R13–DD WT (residues 2300–2541) is a constitutive dimer, consistent with our previous study^7^. In contrast the R13–DD L2509P mutant runs as a monomer, confirming that the proline is disrupting talin dimerisation (Fig.3D).

### Most of the talin mutations have only minor effects on cell morphology, except for the L2509P mutation

We transiently transfected talin double knock–out (TLN1^−/−^TLN2^−/−^) mouse kidney fibroblasts (MKF)^36^ with full length talin–1 constructs containing the mutations shown in Table 1 using equal quantities of plasmid. Western blot analysis ensured translation of full–length proteins (Fig.S3A,E). Whereas, in the biochemical section above, the I392N and L1539P were hard to work with due to difficulties in generating sufficient pure, intact mutant domains, this issue was not evident in the full length mammalian expression, with all the twelve constructs expressing full length talin protein. However, we observed decreased protein expression level for I392N and slightly increased expression for S1750F and D2086V, but otherwise the expression levels were constant (Fig.S3A,E). The adhesion size and abundance were analysed from the cell periphery and did not reveal significant changes in either the adhesion area or adhesion number with the various talin mutations (Fig.S4B,C).

To evaluate the effect the mutations have on cell morphology, we visualised talin and vinculin in transfected fibroblast cells devoid of talins (Fig.4A) and quantified the effect of the mutations on cell area and circularity (Fig.S4D,E). The mutations caused little variance on cell morphology when compared to WT, except for cells carrying the dimerisation domain (DD) mutation L2509P, which were significantly smaller than the WT expressing cells and showed a more circular cell phenotype (Fig.S4D,E), indicating a loss in cell polarisation.

**Figure 4:**
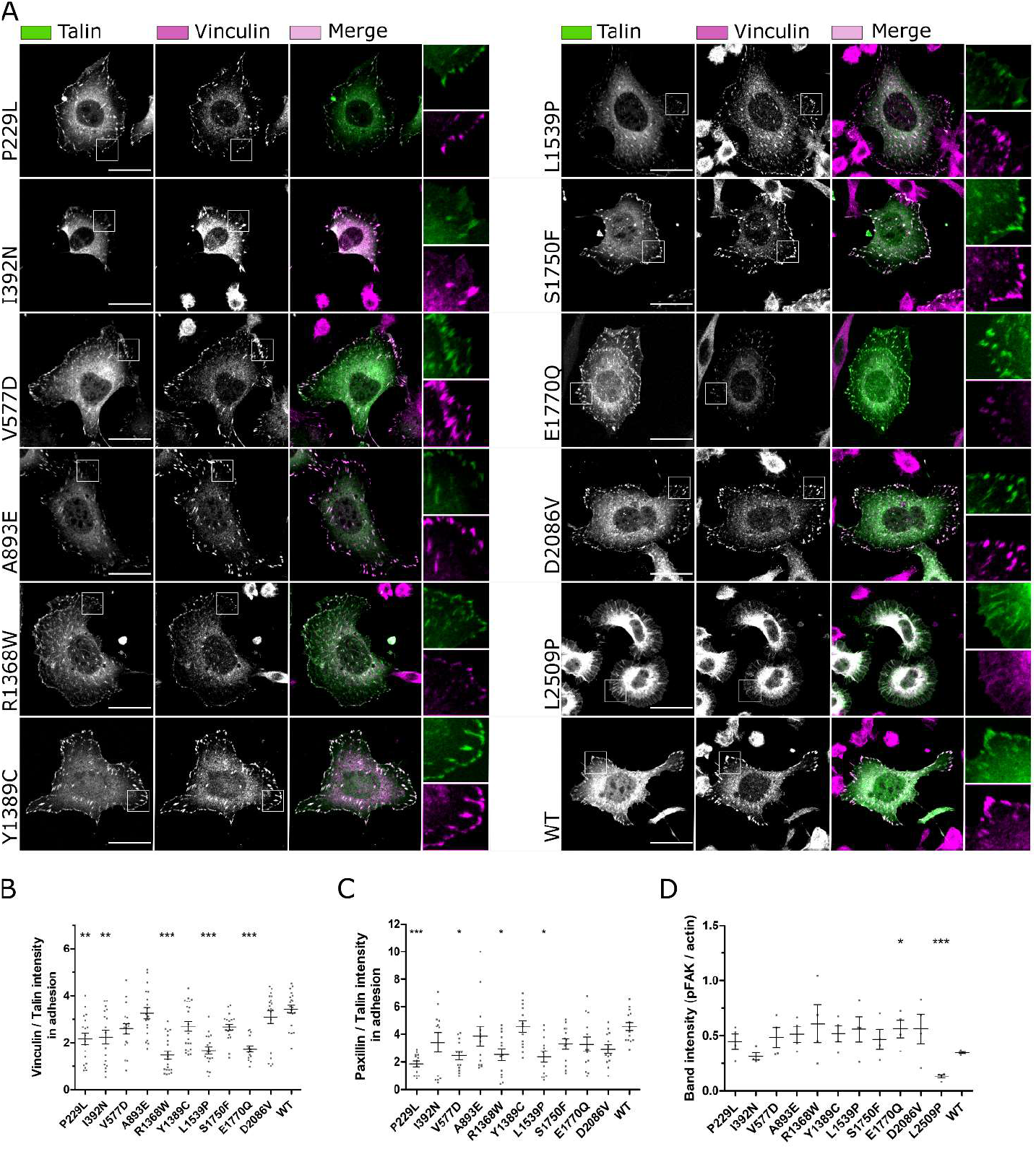
Talin mutations influence the colocalization with vinculin, paxillin and pFAK. A) SUM projections of z–stacks of TLN1^−/−^TLN2^−/−^ mouse kidney fibroblast cells expressing GFP–tagged talin–1 forms and immunolabeled for vinculin. Scale bars are 25 μm, zoom–in square size is 12.5 μm x 12.5 μm. B-C) Analysis of vinculin (B) and paxillin (C) colocalisation with talin in adhesions; n~20 cells per mutation from two separate experiments. D) FAKpTyr397 expression levels quantified from four western blots. The statistical significance in (B) and (C) was analysed by one–way ANOVA and Bonferroni test: *P<0.05, **P<0.01, ***P<0.001. The statistical analysis in (D) was calculated by unpaired t-test.

We next assessed the levels of vinculin, and paxillin within the talin–1 rich adhesion sites with the aid of immunofluorescence labelling and confocal imaging (Fig.4B,C), and assessed the total level of phosphorylated FAK in cells via western blotting (Fig.4D;Fig.S3F). FAK phosphorylated at tyrosine 397 (FAKpTyr397) is a marker for adhesion maturation and corresponds with mechanical activation of talin. Very little FAKpTyr397 is present in non–transfected talin double knock–out MKF cells^37^, which enabled us to monitor the relative levels of FAK phosphorylation with western blotting. Interestingly, several mutants showed decreased levels of vinculin and paxillin within the adhesion sites compared to cells expressing WT talin–1, with the mutants R1368W, L1539P and E1770Q showing significantly less recruitment (Fig.4B,C). We noticed that total vinculin expression level was not majorly influenced by the expression of the talin–1 mutants (Fig.4A;Fig.S3C). In contrast, expression of the talin mutants, P229L, V577D, A893E, L1539P and D2086V led to lower levels of the total paxillin expression when compared to WT–talin expressing cells (Fig.S3D). FAKpTyr397 levels were constant with all mutants except a small increase seen with E1770Q and drastic decrease with L2509P when compared to WT talin expressing cells. With L2509P there was also a decrease seen in total FAK levels (Fig.S3B,G) suggesting that the lack of adhesion maturation and cell polarisation can affect FAK levels in general.

In order to study the L2509P mutation in more detail, we engineered a series of truncated talin–1 constructs with deletions in the c–terminus as follows: WT (residues 1–2541), ΔDD (residues 1–2493) and ΔR13–DD (residues 1–2299) (Fig.5A). Immunofluorescence analysis of the cells transfected with L2509P, ΔDD and ΔR13–DD all showed the same cell morphology (Fig.S4F,G) accompanied with the loss of; i) adhesion maturation, ii) localisation of the FA components paxillin and vinculin, and iii) filamentous actin within he adhesion sites, indicating that the L2509P point mutation disrupts dimerisation and interaction with actin to the same extent as deletion of the entire domain (Fig.5B;Fig.S4H).

**Figure 5.**
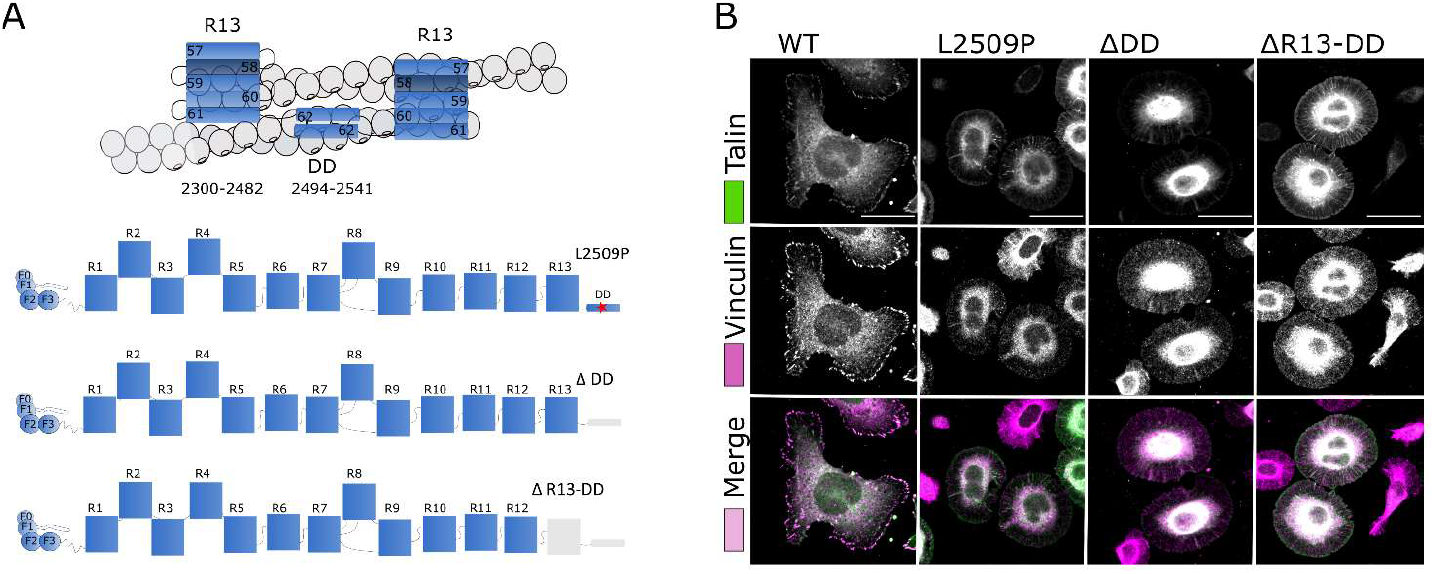
The point mutation L2509P has the same effect on cell morphology as the deletion of the whole dimerisation domain. A) Cartoon of the R13–DD bound to actin (top) and schematic representations of the point mutation L2509P in full–length talin and the truncations; ΔDD and ΔR13–DD. B) SUM projections of z–stacks of TLN1^−/−^TLN2^−/−^ mouse kidney fibroblast cells expressing WT, L2509P, ΔDD, ΔR13–DD talin and immunolabeled against vinculin. No clear localisation of vinculin was evident with any of the mutants. Scale bars are 25 μm.

### Predicting the effect of mutations on talin interactions with other adhesion proteins through cellular and biochemical analysis

The recombinant R7–R8 WT and mutant (R1368W and Y1389C) proteins were used to assess interactions with ligands using biochemical assays. Two of the vinculin binding sites of talin are located in the R7–R8 region, and their exposure is regulated by stability of the α–helical bundles in response to mechanical load^38–40^. To determine whether vinculin Vd1 domain (residues 1–258) binding to talin was affected by the R7–R8 R1368W and R7–R8 Y1389C mutations, we used a size exclusion chromatography (SEC) assay as described previously^38^. Our previous study showed that despite containing two VBS, only one of them is accessible to vinculin^38^, and here we found that R7–R8 WT and R7–R8 R1368W both only bind one vinculin Vd1 molecule (Fig.6A,B). In contrast, the R7–R8 Y1389C mutation, which is in the core of the R7 domain, can bind two Vd1 molecules (Fig.6C), indicating that this mutation enhances the accessibility of the R7 vinculin binding site and alters the stoichiometry of the R7–R8–vinculin interaction. On its own the R7–R8 Y1389C mutation runs as a monomer (Fig.3A; Fig.6C), eluting at the same position as the R7–R8 WT. So, the 2:1 vinculin stoichiometry is due to the R7 domain being destabilised making the VBS more accessible rather than the protein being denatured.

**Figure 6:**
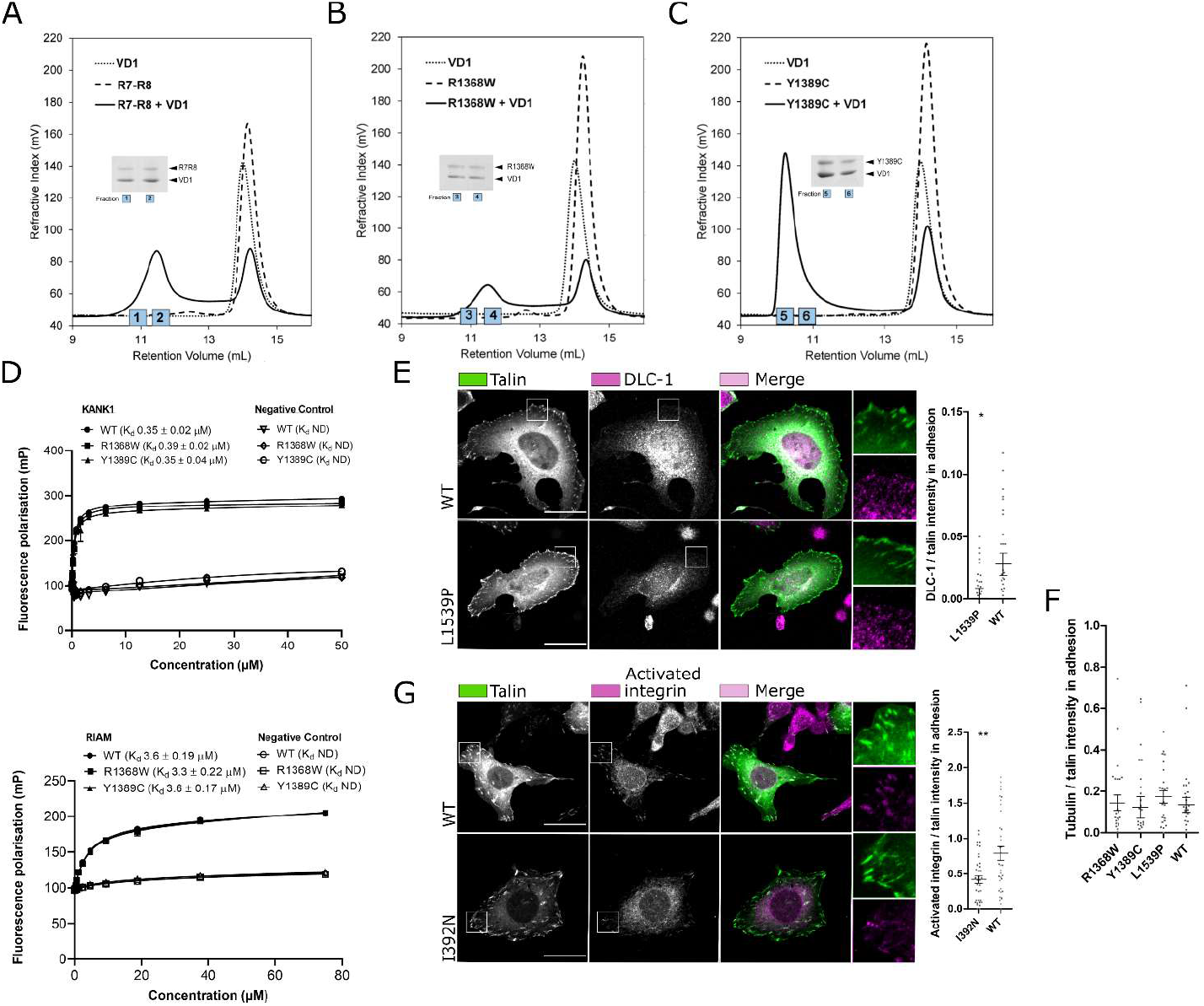
Mutations alter the interaction and colocalisation of talin with vinculin, DLC–1 and activated β1– integrin. A–C) Vinculin Vd1 binding analysed by size exclusion chromatography for R7–R8 WT (A), R7–R8 R1368W (B) and R7–R8 Y1389C (C) purified as recombinant proteins in *E. coli*. The SDS–PAGE gel of the elution fractions is shown. D) Fluorescence Polarisation (FP) assay for purified R7–R8 WT/mutant binding to KANK (top) and RIAM (bottom) peptides. R7–R8 R1368W and R7–R8 Y1389C showed no significant changes in the interaction with KANK and RIAM compared with R7–R8 WT. Fluorescence polarisation assays were performed using protein serially diluted from a starting concentration of 60 μM with target KANK1 (30–68) peptide at 1 μM and 75 μM with target RIAM (4– 30) peptide concentration at 1 μM. Measurements were taken using a CLARIOstar plate reader (BMGLabTech) at 20°C. GraphPad Prism 7 software was used for data analysis with one–site total binding equation used to generate a Kd. E) Representative confocal immunofluorescence images of the co–localisation of DLC–1 in the TLN1^−/−^TLN2^−/−^ mouse kidney fibroblast cells transfected with full length talin WT and L1539P. F) Data obtained from the colocalisation of tubulin in adhesion sites. G) Representative confocal immunofluorescence images of the co–localisation of activated integrin CD29 organisation in the cells transfected with full length talin WT and I392N. SUM projections of z–stacks of cells expressing GFP–tagged talin–1 (WT and/or point mutated) and immunolabeled against integrin CD29 or DLC–1, and ~30 cells per label have been analysed. The statistical significance of all results was analysed by one–way ANOVA and Bonferroni test: *P<0.05, **P<0.01, ***P<0.001. Scale bars are 25 μm, zoom–in square size is 12.5 μm x 12.5 μm.

Both R7 and R8 contain binding sites for proteins that contain LD–motifs and so a Fluorescence Polarisation (FP) assay was used to determine the impact of these mutations on binding of the LD–motifs of KANK1 and RIAM to R7 and R8 respectively. Neither KANK1 binding to R7 nor RIAM binding to R8 had significant differences in binding affinity with the R7–R8 R1368W and R7–R8 Y1389C mutants as compared to R7–R8 WT confirming that these mutations do not perturb the LD–motif binding surfaces (Fig.6D). The talin–KANK1 interaction is important for coordinating the targeting of microtubules to adhesion sites^9^ and consistent with the negligible impact on the talin-KANK1 interaction, no apparent changes in tubulin organisation between WT, R1368W, Y1389C, and L1539P were observed in cell experiment (Fig.6F; Fig.S5A).

Since L1539P is located near to the DLC–1 binding site we also assessed the mutations effect on DLC–1 recruitment to the adhesions in mouse fibroblasts. For this we quantified the amount of DLC–1 within the adhesions and found that L1539P leads to ~42% decrease in DLC–1 amount within the adhesion as compared to the WT (Fig.6E). As I392N is in close proximity to the integrin binding site in F3, (PDB:2H7D)^41^, we assessed whether the mutation can affect the talin–mediated integrin activation. For this, we quantified the amount of activated integrins within adhesions. Expression of I392N lead to ~53% decrease in the amount of activated integrin when compared to WT talin (Fig.6G).

### Cancer–derived talin–1 point mutations affect cell migration, invasion and proliferation

To study the functional characteristics of the talin mutants, we measured random cell migration speed on cells cultured on fibronectin–coated coverslips (Fig.7A,B). WT–transfected cells had an average migration speed of 0.65 μm/min. I392N was the only mutant that caused increased migration speed (0.82 μm/min). In contrast, R1368W caused a slight decrease and L2509P caused a marked decrease in cell migration with average speeds of 0.54 and 0.37 μm/min respectively, with the migration speed of L2509P close to that of non–transfected cells (0.35 μm/min) (Fig.S5B).

**Figure 7:**
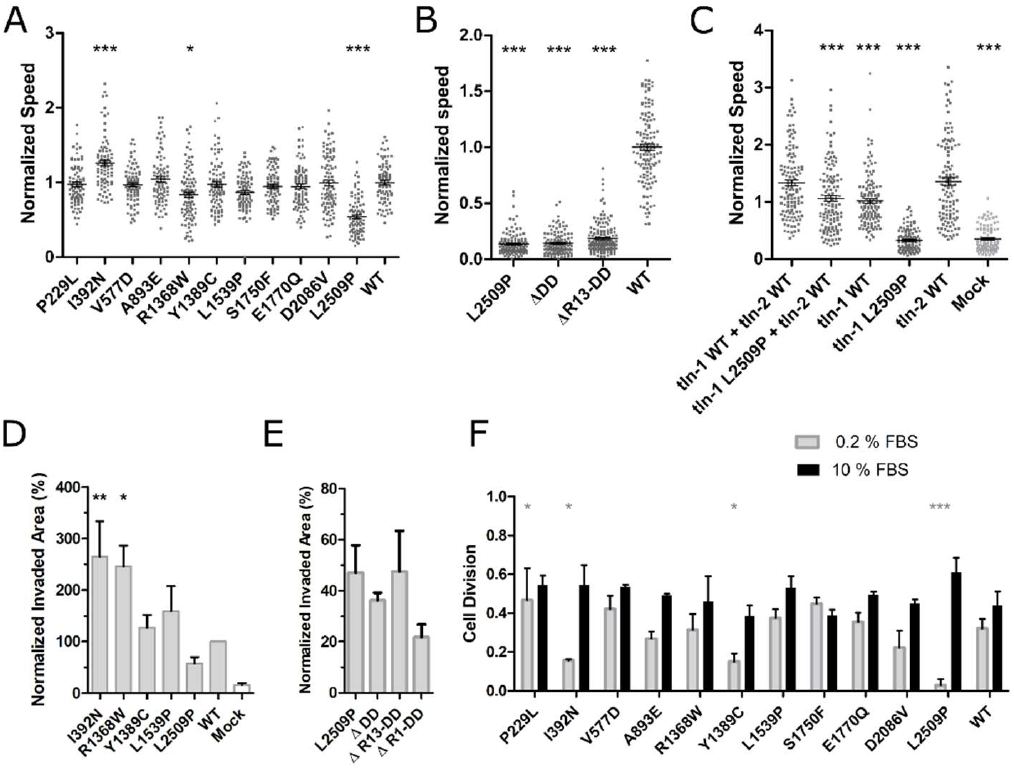
Talin mutations affect cell migration and proliferation. A) Random migration speed (μm/min) determined from time–lapse images of talin expressing cells. B) Migration speed on 2D surface showing reduced migration speed in all mutated/truncated constructs in comparison to WT. C) Migration assay on 2D surface showing reduced migration speed in cells co–transfected with talin–1 L2509P and full–length WT talin–2. The statistical significance of all results analysed in comparison to talin–1 WT + talin–2 WT by one-way ANOVA and Bonferroni test: *P<0.05, **P<0.01, ***P<0.001. The results are normalised with talin–1 WT. D) Invasion assay through Matrigel matrix towards 10% FBS containing medium. Control cells were mock–transfected with GFP–expressing plasmid. The values are normalised to WT and statistical significance measured in comparison to WT. Data are mean +/−SEM. The statistical significance was analysed by one–way ANOVA and Bonferroni test: *P<0.05, **P<0.01, ***P<0.001. E) Cell invasion through Matrigel in 3D environment showing the invasiveness potential of the L2509P and truncated talin constructs. Invasion assay was repeated at least three times in triplicate chamber for each construct on separate days. F) Cell proliferation analysis in the presence of 10% FBS and 0.2% FBS, showing the number of times the cells divide in 12 h; n~80 cells per mutation from four separate experiments. The statistical analysis was calculated by t–test, non–parametric test of Mann–Whitney: *P<0.05, **P<0.01, ***P<0.001 compared to WT for each condition.

Since many cell types express both, talin–1 and talin–2, we were curious to see whether the L2509P mutation in talin–1 would show a phenotype when expressed together with talin–2. Indeed, we found that the mutation caused significant decrease in migration speed in cells co–transfected with talin–2 when compared to cells expressing both WT talins (Fig.7C).

Based on the migration assay, we selected I392N, R1368W, Y1389C, L1539P and L2509P for further analysis. We characterised how these five mutants affected the ability of cells to invade through a 3D Matrigel matrix (Fig.7D). I392N and R1368W showed the highest invasion rates, whereas Y1389C and L1539P did not significantly differ from WT. Further, the mutant L2509P, which showed a poorly polarised cell phenotype (Fig.5B) and significantly reduced migration speed on 2D culture (Fig.7A,B), was only able to promote limited cell invasion in 3D (Fig.7D). The impact on cell migration and invasion, was the same for the L2509P and the truncated mutants whether we removed the entire rod, R13–DD, DD or applied the point mutation L2509P (Fig.7B,E).

In full–serum conditions the mutants and the mock–transfected cells (Fig.S5C) showed equally efficient cell proliferation rate when compared to WT talin–1 expressing cells (Fig.7F). However, during serum depletion, I392N, Y1389C, L2509P and mock–transfected cells showed significant decrease (~53%, ~53%, ~91% and ~45%, respectively) and the P229L showed ~30% increase in cell division as compared to the cells transfected with WT talin–1 (Fig.7F; Fig. S5C), suggesting that talin–1 mediated changes can affect cell proliferation.

## Discussion

Loss of anchorage–dependent growth, changes in ECM remodelling, and cytoskeletal changes are necessary for cancer progression^42^. Talins are central regulators of these processes, providing mechanical linkages between ECM and the actin cytoskeleton, In this study we explored whether mutations in talin identified in large scaled cancer exome sequencing might contribute to cancer progression.

There are two isoforms of talin, talin–1 and talin–2, which show different expression patterns^43, 44^ While talin–1 is expressed in all tissues, talin–2 has more variability and generally lower expression levels. Overall expression levels, derived from GEPIA (gene expression profiling interactive analysis) server^45^, indicate that talin–1 expression levels are more variable between different cancers and their counterpart healthy tissues than those of talin-2 (Fig.S6) with several cancer types showing significant changes in talin–1 expression level. The highest overexpression is associated with Glioblastomas (GBM; >300%), Brain Gliomas (LGG; >300%) and Pancreatic adenocarcinomas (PAAD; >300%). In contrast, the most drastic talin–1 downregulation is seen in uveal melanoma (UCS; −25%) and endometrial carcinoma of the uterine (UCEC; −25%).

These datasets are large and expansive, and so we prioritise bioinformatic analysis as the first stage in our pipeline approach enabling us to explore the large quantity of talin–1 mutations and predict the impact of the mutations on the stability of protein structure. With the aid of bioinformatic classification, molecular modelling, biochemical analyses and functional cell biology assays, we found that mutations in talin–1 affect cellular processes linked with cancer progression, such as migration, invasion and proliferation (Table 2). By this approach, we identified mutations located in the previously identified binding sites showing strong destabilization of the domain in free energy calculations (I392N, R1368W, Y1389C, L1539P and L2509P) for further investigation. Several of the studied mutations were difficult to produce for the biochemical assays, suggesting that these indeed destabilised the corresponding talin domain. However, all of the mutations could be efficiently produced within cells as full-length proteins that were localised to the adhesions. Mutation within the dimerization domain, L2509P, was the only mutation leading to disruption of the adhesion complexes, due to lack of dimerization and actin binding. In contrast the other mutations showed more subtle changes in migration and invasion or in amount or activity of the binding partners within adhesions. Mutations I392N, Y1389C and L1539P had the highest initial bioinformatics scores and this correlated well with the data gained from cell experiments. We showed increased migration and invasion and changes within integrin activation status for cells expressing I392N. For Y1389C and L1539P we observed changes in vinculin binding and DLC1 recruitment to the adhesions, respectively. Mutation R1368W led to changes in migration, invasion and in recruitment of binding partners although it received an intermediate score in the bioinformatics analysis. Finally, while L2509P received only moderately high score in bioinformatic evaluation, it was found to be highly pathogenic, causing abolishment of talin dimerization.

**Table 2.**
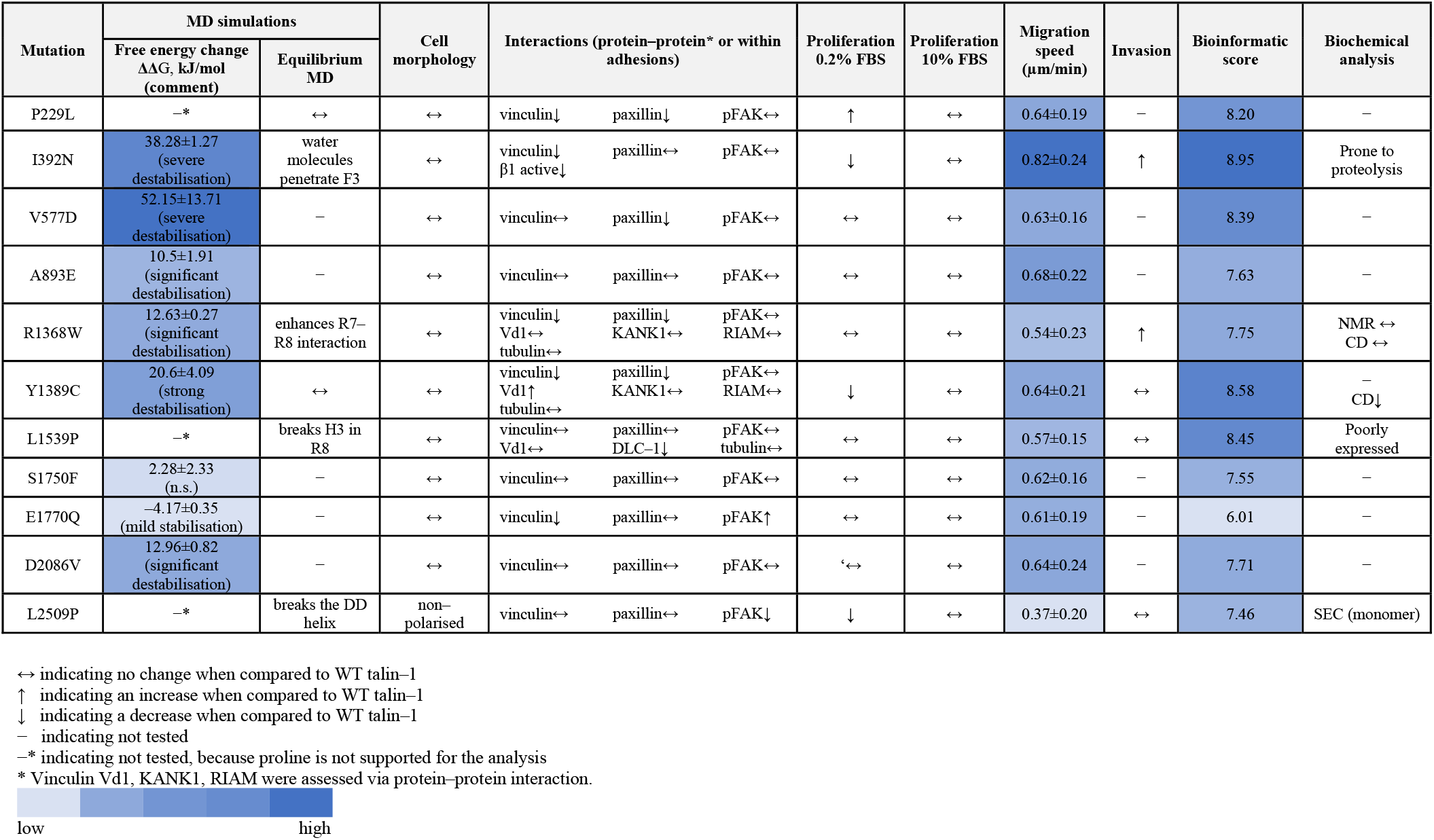
Summary table of the analysis of talin–1 mutations. Migration speed for the WT–talin was 0.65±0.18 μm/min.

Out of the eleven mutants studied here, I392N, originally found from pancreatic carcinoma, had the most pronounced effect in terms of driving migration and invasion. A previous study by Isenberg et al. proposed the amino acids 385–406 of F3 helix as a potential membrane–anchoring domain with I392 as one of the residues that inserts into the lipid bilayer^30^ and mediates integrin activation^46^. Our results showed a clear decrease in the amount of active integrin in talin–rich adhesions, when compared to WT (Fig.6G).The mutated helix in F3 experiences mechanical tension applied on talin^47^ and this might result in increased dissociation of the destabilised mutated helix from the rest of the F3 under applied load, influencing integrin association and thereby contributing to the adhesion turnover/dynamics. Biochemical experiments with this mutant were hampered by proteolytic truncation of the mutated talin head during recombinant protein expression, indicating folding defect in the C–terminal part of the protein.

The R7–R8 domains have cooperative function, with R8 sitting outside the force transduction pathway, and protected from mechanical stress by R7^40^. In addition, there are binding sites for multiple molecules such as KANK1, DLC–1, RIAM, paxillin and vinculin in both domains. One surprising result from this study was the observation that in every MD simulation carried out for R7– R8 (WT, R1368W, Y1389C, and L1539P), the R7 and R8 subdomains were found to transiently interact over the course of the 100 ns MD. Furthermore, the MD simulations suggested that the R1368W mutant in R7 might enhance these interactions between R7 and R8 domains (Fig.2B;Fig.S2A,B). All the crystal structures of talin R7–R8 reported to date have shown an open conformation where there are limited contacts between the R7 and R8 subdomains^11, 38, 48^ and such an interaction was not readily apparent in the NMR analysis (Fig.S4A). However, the recent crystal structure of TLNRD1 (talin rod domain containing protein 1 (PDB:6XZ4^49^), a protein structurally homologous to talin R7–R8, revealed a similar close association of the two subdomains, supporting the notion that the two talin subdomains might also interact. One possibility is that the interaction between two domains might reduce accessibility to ligand binding sites on the domains, and perturb the localisation of the FA markers within adhesions (Fig.4B,C), affecting cell behaviour. However, binding affinity of the R1368W mutant for KANK1 and protein stability remained consistent with WT, suggesting that the cellular effects we observed are not due to alterations in known functions of R7. It is possible that this mutation is altering the dynamics of the R7 and R8 domains causing the altered cellular behaviour we observe. Therefore, the role of such an interaction is not fully clear and warrants further investigation.

The VBS in R7 is one of the hardest to expose^38^ but can be stretch activated^40^, interestingly, the R7–R8 Y1389C mutation significantly enhanced the VBS accessibility. Gel filtration analysis of vinculin Vd1 binding to R7–R8 Y1389C revealed that while R7–R8 WT and the R7–R8 R1368W mutant only bind a single Vd1 molecule, the R7–R8 Y1389C was able to bind two Vd1 (Fig.6A,B,C). This suggests that introduction of the mutation destabilises the R7 helical bundle, allowing vinculin binding more readily in the absence of force. Furthermore, reduced R7 stability will likely have a knock–on effect on R8 stability which may indirectly lead to signalling defects by perturbing R8^12^. The additional actin recruitment via vinculin could have a direct effect on FA dynamics by facilitating formation of talin–vinculin pre–complexes, necessary to enable efficient adhesion maturation^50^. This could also affect actomyosin contractility and Rho/ROCK signalling, which have been shown to regulate cell proliferation^51^.

DLC–1 is a tumour suppressor^52^ and contains an LD–like motif, which is required for the full tumour suppressor activity of DLC–1 via interaction with talin^46^. DLC–1 residue D470 makes direct contact with the positively charged side chain of K1530 and K1544 in R8^11^. In cells transfected with L1539P mutant, the colocalisation between talin and DLC–1 was significantly decreased (Fig.6E), indicating that this mutation interferes with DLC-1 binding.

In previous studies, talin depletion caused a halt in cell cycle of epithelial cells that could be rescued with expression of C–terminal talin that was able to cause FAK phosphorylation^53^. The linkage between the C–terminal actin–binding site, ABS3 and actin is required for polarisation of cells^37^ and this linkage depends on talin dimerisation. Mutation in the dimerization domain, L2509P, resulted in an unpolarised and non–migratory cell phenotype lacking mature FAs and actin contacts on the cell surface. This mutation sits in the talin dimerisation domain, a single helix which forms an antiparallel dimer^7, 54^. The same phenotype was observed with complete removal of the dimerisation domain helix (ΔDD) as with L2509P point mutation. Our biochemical results showed that the R13–DD L2509P mutant resulted in complete loss of dimerisation compared to the R13–DD WT which is a constitutive dimer in solution (Fig.3D).

In this study, we systematically explored hundreds of talin–associated mutations and the bioinformatics pipeline described here enabled us to rapidly and robustly screen this huge number and narrow it down to an experimentally tractable subset for detailed analysis. Using this approach, we selected 11 mutants for thorough analysis and found that the bioinformatic scores reflected the corresponding cell phenotype well in most cases, however, the pathogenicity of the mutations in tissue and cell context is difficult to assess. Here the cell phenotype of cancer–associated talin mutations was observed without the effect of the other mutations commonly found in cancer cells. As the mutations within the common oncogenes will heavily affect the cell signalling, it is difficult to predict exactly what would be the contribution of these studied talin–1 mutations for the cancer progression in their original context. It is however noteworthy that several talin point mutations influenced cell migration, invasion and cell polarisation even when expressed without the other cancer–associated factors.

Investigation of the contribution of talin to cancer progression is timely. During this study ~670 more cancer–associated talin–1 point mutations have been added to the COSMIC database (Fig.S1) and there are recent studies discussing the connection between talin and cancer^23, 55–58^. The work we present here demonstrates an efficient and fast pipeline approach using bioinformatics tools to characterise future talin mutations identified in cancer and other diseases.

## Materials and methods

### Cell lines and talin constructs

Theodosiou et al.^36^ previously described the TLN1^−/−^TLN2^−/−^ mouse kidney fibroblast (MKF) cell line. Cells were maintained in a humidified 37°C, 5% CO_2_ incubator. High glucose Dulbecco’s modified Eagle medium (DMEM) supplemented with 10% fetal bovine serum (FBS) was used in all experiments except in the starvation conditions where 0.2% serum was used. The cell line was regularly tested for mycoplasma contamination. Talin variants were subcloned into a modified pEGFP–C1 vector backbone (Clontech). Cells were transfected with 6 μg plasmid DNA per 10^6^ cells using Neon transfection system (Thermo Fisher Scientific) using parameters 1400 V, 30 ms, one pulse. The expression constructs for cell culture experiments with the c–terminal EGFP–tag are as follows: wildtype talin–1 (1–2541); ΔR13–DD (1–2299); ΔDD (1–2493); ΔR1–DD (1–481) and the point mutants in the full–length talin–1 P229L, I392N, V557D, A893E, R1368W, Y1389C, L1539P, S1750F, E1770Q, D2086V, L2509P.

### Migration and Matrigel invasion analysis

Transfected cells were incubated for 24 h, trypsinised and plated on well–plates coated with 10 μg/ml fibronectin. Cells were allowed to attach for 90 minutes, after which the media was changed. The time–lapse images captured with EVOS FL auto microscope (Thermo Fisher Scientific) were analysed manually using ImageJ (Fiji) and MTrackJ plugin^59, 60^.

Corning BioCoat Matrigel Invasion Chamber containing an 8–micron pore size PET membrane with a thin layer of Matrigel basement membrane matrix were used for the invasion assay. Transfected TLN1^−/−^TLN2^−/−^ MKF cells were cultured overnight, followed by cultivation in starvation medium containing 0.2% FBS for 40–45 h. Number of transfected cells was measured by Luna–FL dual Fluorescence Cell Counter (BioCat GmbH) Chambers prepared according to the manufacturer. DMEM medium containing 10% FBS was used as chemoattractant in the lower level of chamber. The chamber plate was incubated at humidified 37°C and 5% CO_2_ incubator for 24 h, after which the cells were fixed with 100% methanol. Cells were stained with 0.2% crystal blue for 10 minutes following by rinsing the excess stain. The non–invaded cells were removed from the upper membrane surface using cotton tipped swab. The inserts were allowed to air dry overnight. The membrane was removed using scalpel and placed bottom side down on a microscope objective slide on which a small drop of immersion oil. The membranes were scanned using PRIOR OpenStand microscope using 20x objective and Jilab SlideStrider software (1.2.0). The invaded cell area was calculated using ImageJ (Fiji). Invasion assay was repeated at least three times in triplicate chamber for each selected construct.

### Immunostaining and confocal imaging

After 24 h transfection, cells were trypsinised and plated on coverslips coated with 10 μg/ml fibronectin and incubated for 24 h. Cells were fixed with 4% paraformaldehyde, permeabilised and immunostained using standard protocol. Antibodies are listed in Table S1.

Immunostained samples were imaged with Zeiss Cell ObserverZ1 inverted microscope and LSM 780 confocal unit (Zeiss, Oberkochen, Germany) using 63x/1.4, WD 0.19 mm oil immersion objective. Images were taken using Zeiss Zen Black software and analysed by ImageJ as described previously^37^. Within each experiment, the imaging parameters were kept constant to allow quantitative image analysis. Detailed image analysis is described in Supplementary material.

### Western blotting

Transfected cells were grown for 24 h, lysed with RIPA buffer supplemented with protease inhibitor cocktail (Sigma–Aldrich lot#126M4016V). After centrifugation, cell lysates were applied on an SDS–PAGE to separate protein. A wet blot system was used to transfer the separated protein from gel onto a polyvinylidene fluoride (PVDF) membrane. Blots were quantified using ImageJ. Antibodies are listed in Table S1.

### Constructs for protein expression in *E. coli*

The talin–1 fragments (head, residues 1–405; R7–R8, residues 1355–1652; R13–DD, residues 2300– 2541), generated using full–length mouse talin–1 as a template, were introduced into a modified pHis vector to create N–terminal His6–tagged constructs. The His6–tag is separated from the talin fragment by an eleven–residue linker: SSSGPSASGTG. Mutagenesis was performed using QuikChange II Site–Directed Mutagenesis kit. Talin constructs were expressed in BL21(DE3) *E. coli* cells and induced with 0.1 mM IPTG at 18°C overnight. Clarified lysates were loaded onto an affinity column (HisTrap HP 5 ml; GE Healthcare). Eluted protein was further purified using an anion exchange column (HiTrap Q HP 5 ml; GE Healthcare) before buffer exchange into PBS and storage at –20°C.

### NMR Spectroscopy and Fluorescence Polarisation Assay

For NMR analysis, talin constructs were grown in 2M9 minimal media with ^15^N–labelled NH4Cl. Protein was purified as above and buffer exchanged into 20 mM Na–phosphate pH 6.5, 50 mM NaCl, 2 mM DTT, 5% (v/v) D2O. NMR spectra were obtained at 298 K on a Bruker AVANCE III 600 MHz spectrometer equipped with CryoProbe. All R7–R8 ^1^H,^15^N-HSQC spectra were obtained at a concentration of 160 μM. For fluorescence polarisation (FP) experiments, mouse KANK1 and RIAM peptides were synthesised by GLBiochem (China) and coupled with either BODIPY or Fluorescein dye via a C–terminal cysteine residue:

KANK1 (30–68) – PYFVETPYGFQLDLDFVKYVDDIQKGNTIKKLNIQKRRK–C
RIAM (4–30) – SEDIDQMFSTLLGEMDLLTQSLGVDT–C

### Size Exclusion Chromatography with Multi–Angle Light Scattering

Talin R13–DD wildtype and R13–DD L2509P were analysed by SEC–MALS at a concentration of 100 μM at room temperature with a Superdex 75 column (GE Healthcare Life Sciences). Eluted proteins were analysed with Viscotek SEC–MALS 9 and Viscotek RI detector VE3580 (Malvern Panalytical). Molecular mass was determined using OmniSEC software. For analysis of Vd1 binding to talin, proteins were incubated at a 2:1 ratio at a concentration of 100 μM and analysed at room temperature.

### MD Simulations

RCSB PDB structures were used as starting conformations for MD: 3IVF for F2–F3 (residues 208–398)^61^, 2H7E for F3 (residues 309–405)^41^, 1SJ8 for R1 (residues 487–656)^62^, 2L7A for R3 (residues 796–909)^63^, 2X0C for R7–R8 and R7 (residues 1352–145 7, 15 85–165 9)^38^, 2KBB for R9 (residues 1655–1822)^33^, 3DYJ for R11 (residues 1975–2140)^64^ and 2QDQ for DD domain (residues 2494–2541)^7^. The F2–F3 and R7–R8 inter-subdomain binding energy was calculated using MM-PBSA^65^. Structural analysis was performed using PyMOL. The secondary structure analysis was based on the Dictionary of Secondary Structure of Proteins (DSSP) algorithm^66^.

Equilibrium MD simulations were performed using Gromacs^67^ at the Sisu supercomputer, CSC, Finland. The CHARMM27 force field^68^ and explicit TIP3P water model^69^ in 0.15 M KCl solution were used. The energy minimisation of the system was performed in 10000 steps. The system was equilibrated in three phases using harmonic position restraints on all heavy atoms of the protein, as described in our previous study^70^. Integration time step of 2 fs was used in all the simulations. The temperature and pressure of the system was maintained at 310 K using the V–rescale algorithm^71^, and 1 bar using Berendsen algorithm^72^. At least three 100 ns replicas were generated for each system.

Alchemical free energy calculations were prepared using PMX^73^ and performed with Gromacs at Puhti supercomputer, CSC, Finland. The Amber99SB*–ILDN force field^74^ and TIP3P water model in 0.15 M NaCl solution were used. Each system was energy minimised for 10000 steps and then equilibrated for 1 ns using harmonic position restraints on all heavy atom of the protein. The temperature and pressure of the system was maintained at 298 K and 1 bar using Berendsen algorithm for the system equilibration, while V–rescale and Parrinello–Rahman^75^ algorithms were used for equilibrium MD and non–equilibrium morphing simulations. Integration time step of 2 fs was used in all the simulations. Each state of the system was run for 100 ns equilibrium MD. 100 non–equilibrium morphing simulations were prepared for each physical state of the system, using snapshots captured from the equilibrium trajectories, linearly spaced from 10.9 to 100 ns. Fast non– equilibrium simulations were morphing the system from one state to another in 100 ps for change conserving mutations, and in 200 ps for charge changing mutations. A soft–core potential^76^ was used for the non–equilibrium simulations. The whole calculation, including system preparation, was repeated three times and average free energy value was obtained. Approximately 5.6 μs MD simulations in total were performed for the analysis. Proline involving mutations were not analysed as proline is not supported for the analysis.

## Acknowledgements

This research was supported by the Academy of Finland (grant 290506 to V.P.H. and grant 323021 to V.V.M.), a Biotechnology and Biological Sciences Research Council grant BB/N007336/1 (B.T.G.) and a Human Frontier Science Program grant RGP00001/2016 (B.T.G.). L.A. received support from the graduate school of Tampere University and the Anu Kirra’s grant foundation. We acknowledge CSC for supercomputing resources and Biocenter Finland for infrastructure support. The authors acknowledge the Biocenter Finland (BF) and Tampere Imaging Facility (TIF) for the services. We thank Prof. Reinhard Fässler and Dr Carsten Grashoff (Max Planck Institute of Biochemistry) for help with TLN^1–/–^TLN^2–/–^ cells; Prof. Michael Sheetz (National University of Singapore) for providing the mouse wildtype talin expression construct. We thank Rolle Rahikainen, Anssi Nurminen and Sampo Kukkurainen (Tampere University) for their support and insights. Ulla Kiiskinen and Niklas Kähkönen (Tampere University) are acknowledged for technical support.

