## Supplementary materials for "Cancer associated talin point mutations disorganise cell adhesion and migration"

**Supplementary Figures**

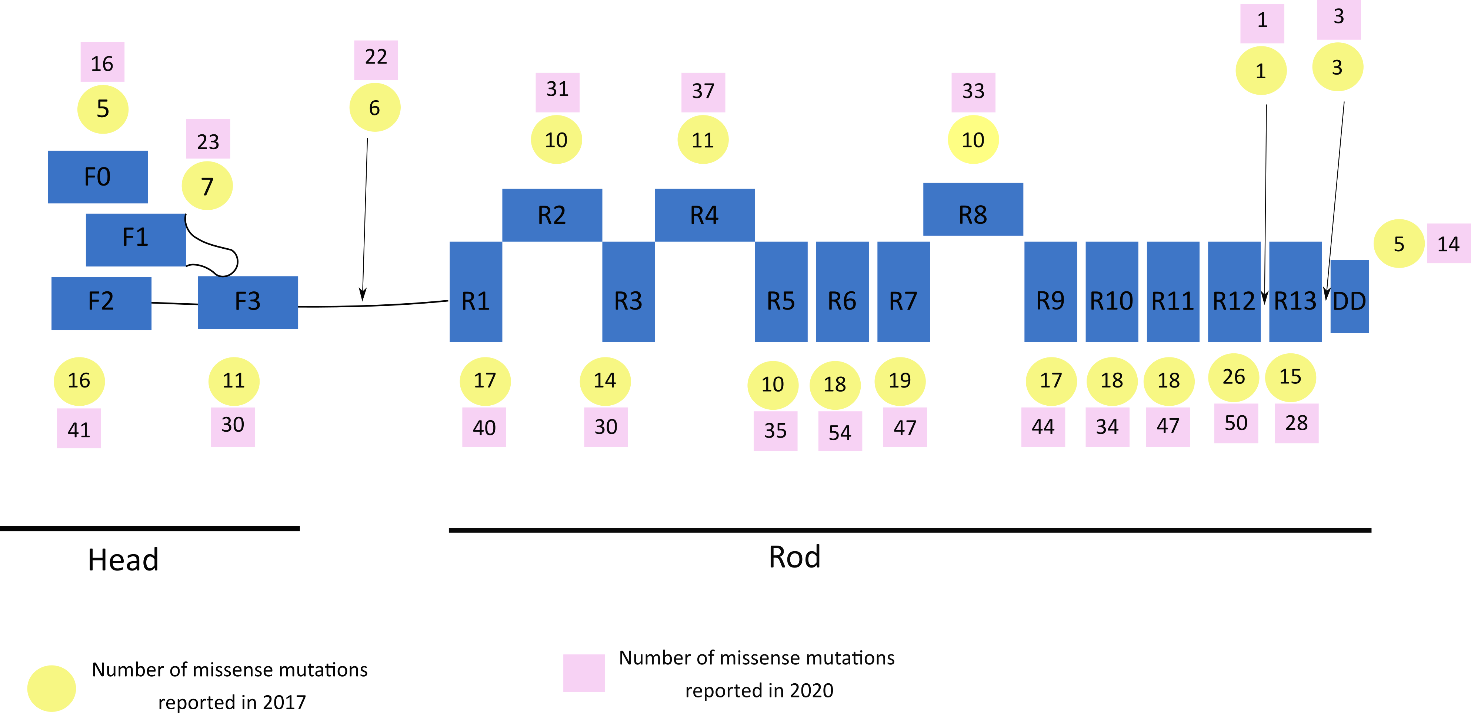

**Figure S1: The number of talin**–**1 missense mutations reported in COSMIC database.** *TLN–1* point mutations investigated from COSMIC database reported in January 2017 (circle) and the list updated in September 2020 (square).

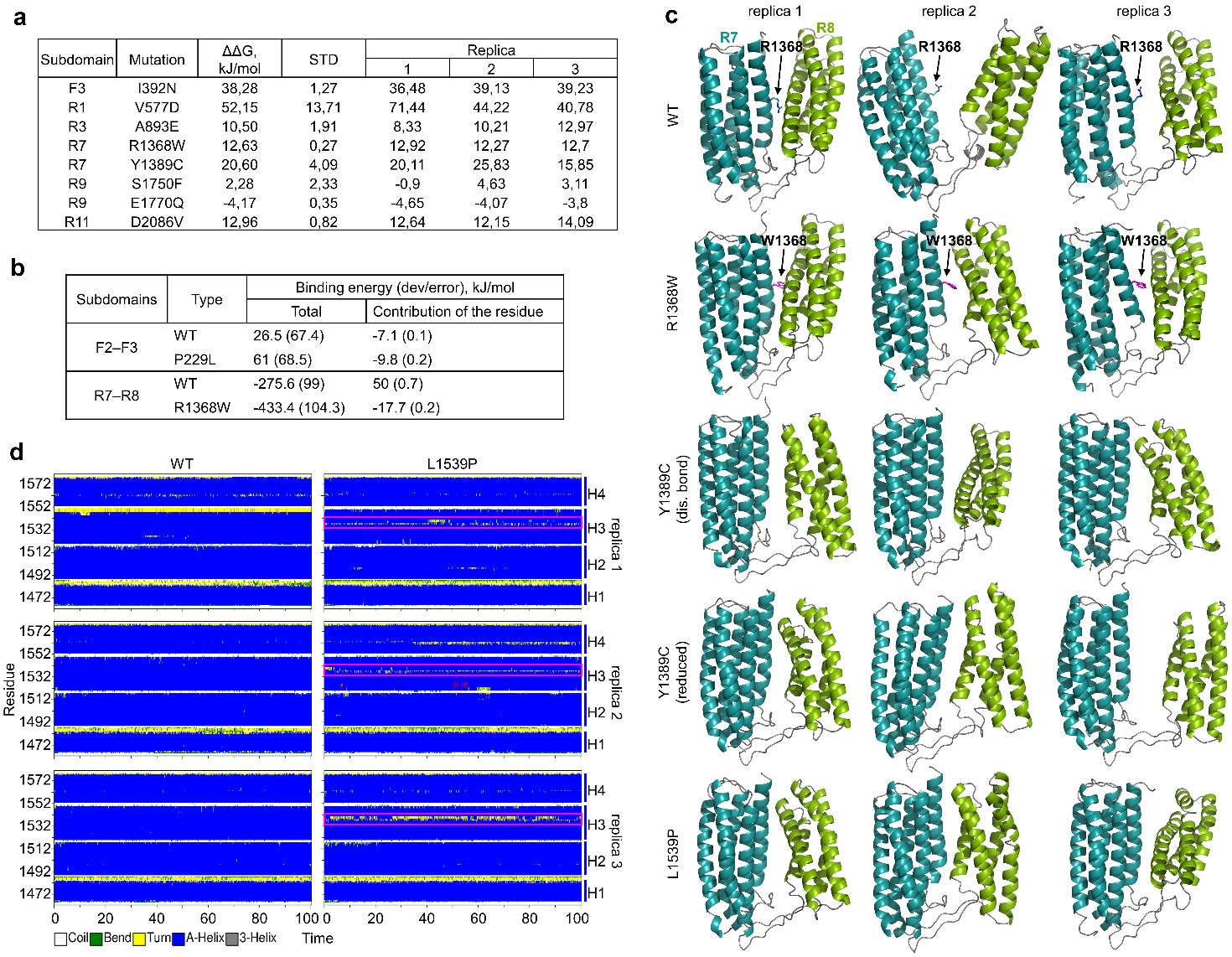

**Figure S2:** (a) Analysis of free energy changes in protein stability upon mutation. Mutations that involve proline were not analysed because proline is not supported for the analysis. (b) MMPBSA analysis of the inter–domain binding energy for F2–F3 WT, F2–F3 P229L, R7–R8 WT and R7–R8 R1368W. (c) Structure snapshots from MD simulations of talin rod R7–R8 fragment for WT, R1368W, Y1389C and L1539P. The snapshots were captured 100 ns. (d) Secondary structure analysis for R8 subdomain in WT and L1539P mutant showing that the mutation breaks H3.

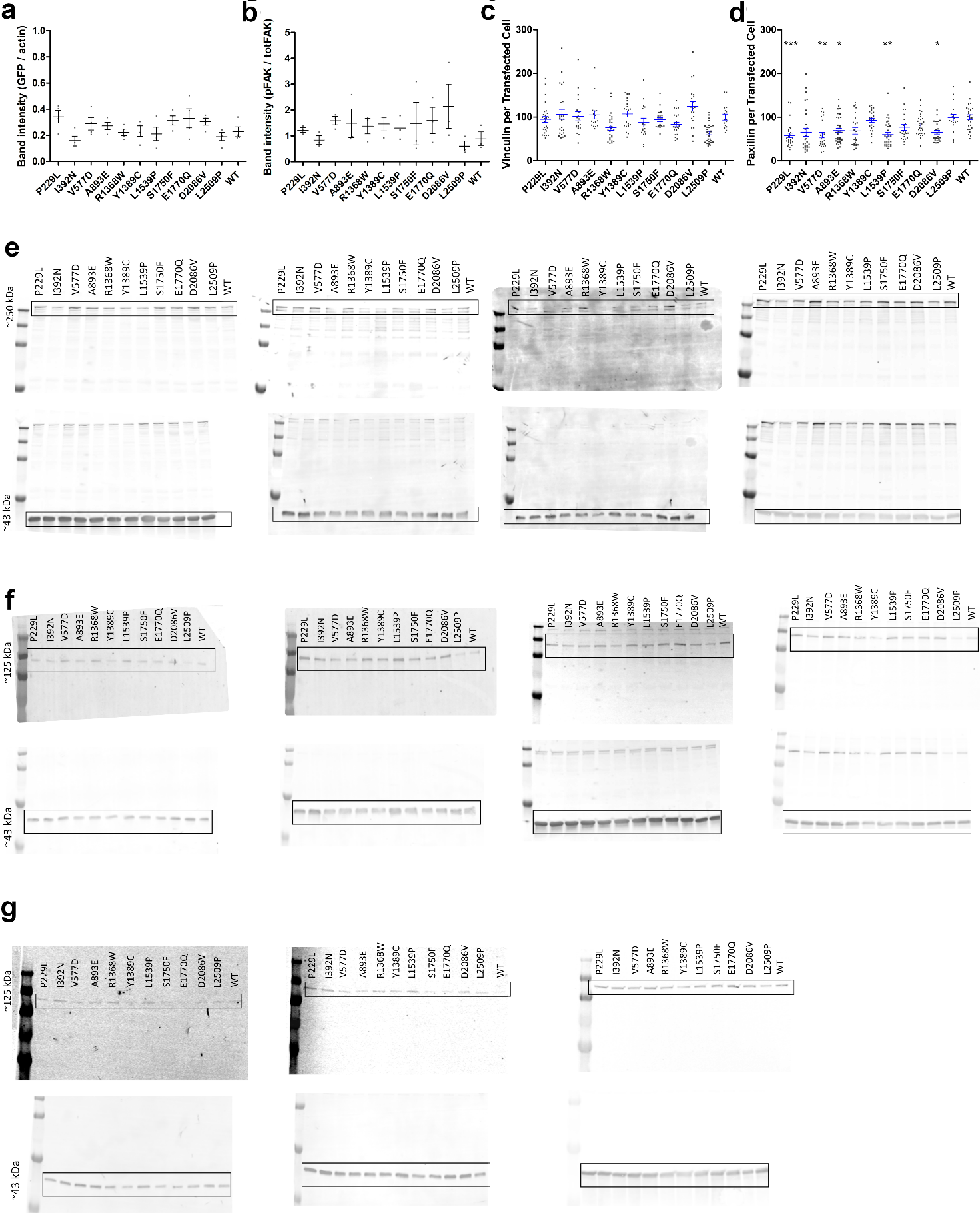

**Figure S3:** (a) Quantification of Western Blots showing talin expression level in the analysed cells (GFP/actin). (b) Quantification of Western Blots showing ratio of pFAK to total FAK expression level in the analysed cells. The statistical analysis in (a) and (b) was calculated by unpaired t–test. (c, d) Total expression levels of immunolabelled vinculin (c), and paxillin (d) quantified from talin expressing cells; n~30 cells per mutation pooled from two separate experiments for each analysis. The statistical significance of all results was analysed by one way ANOVA and Bonferroni test: P>0.05, not significant. (e) Western Blots used for quantification of talin expression. Blots are immunolabelled against GFP (top) and actin (bottom). (f) Western Blots used for quantification of FAKpTyr397. Blots are immunolabelled against FAKpTyr397 (top) and actin (bottom). (g) Western Blots used for quantification of total FAK. Blots are immunolabelled against total FAK (top) and actin (bottom).

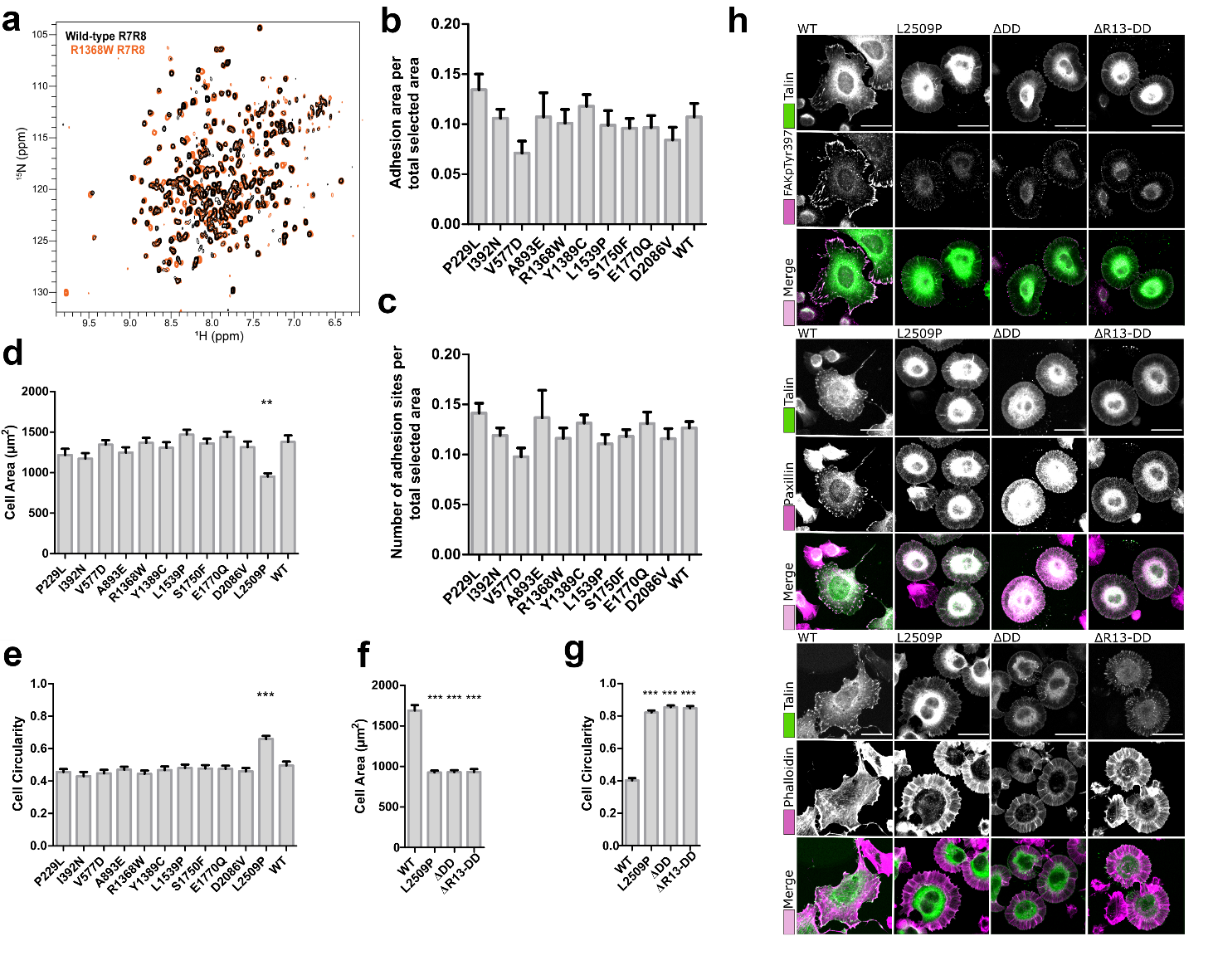

**Figure S4:** (a) NMR spectra of R7–R8 WT (black) and R1368W (orange). (b, c) Graphs showing the results of adhesion site analysis; n~30 cells from two separate experiment. Results are shown as the ratios of adhesion area per total selected area from cell (b) or as number of adhesions per selected area (c). The statistical significance of all results was analysed by one way ANOVA and Bonferroni test: P>0.05, not significant. Due to lack of polarity and seemingly disturbed focal adhesions, L2509P was not included in this analysis. (d, e, f, g) Cell area (d, f) and circularity (e, g) were quantified from microscopy images of cells expressing the transiently transfected talin–1 point mutant constructs; n~40 cells per mutation pooled from four separate experiments. The statistical significance of all results was analysed by one way ANOVA and Bonferroni test: P>0.05, not significant. (h) SUM projections of z–stacks of cells expressing WT, L2509P, ∆DD, ∆R13–DD talin and immunolabeled against FAKpTyr397 and paxillin. Fluorochrome–conjugated phalloidin was used to visualise actin filaments. No clear localisation of any of the FA components was evident with any of the mutants. Scale bars are 25 µm.

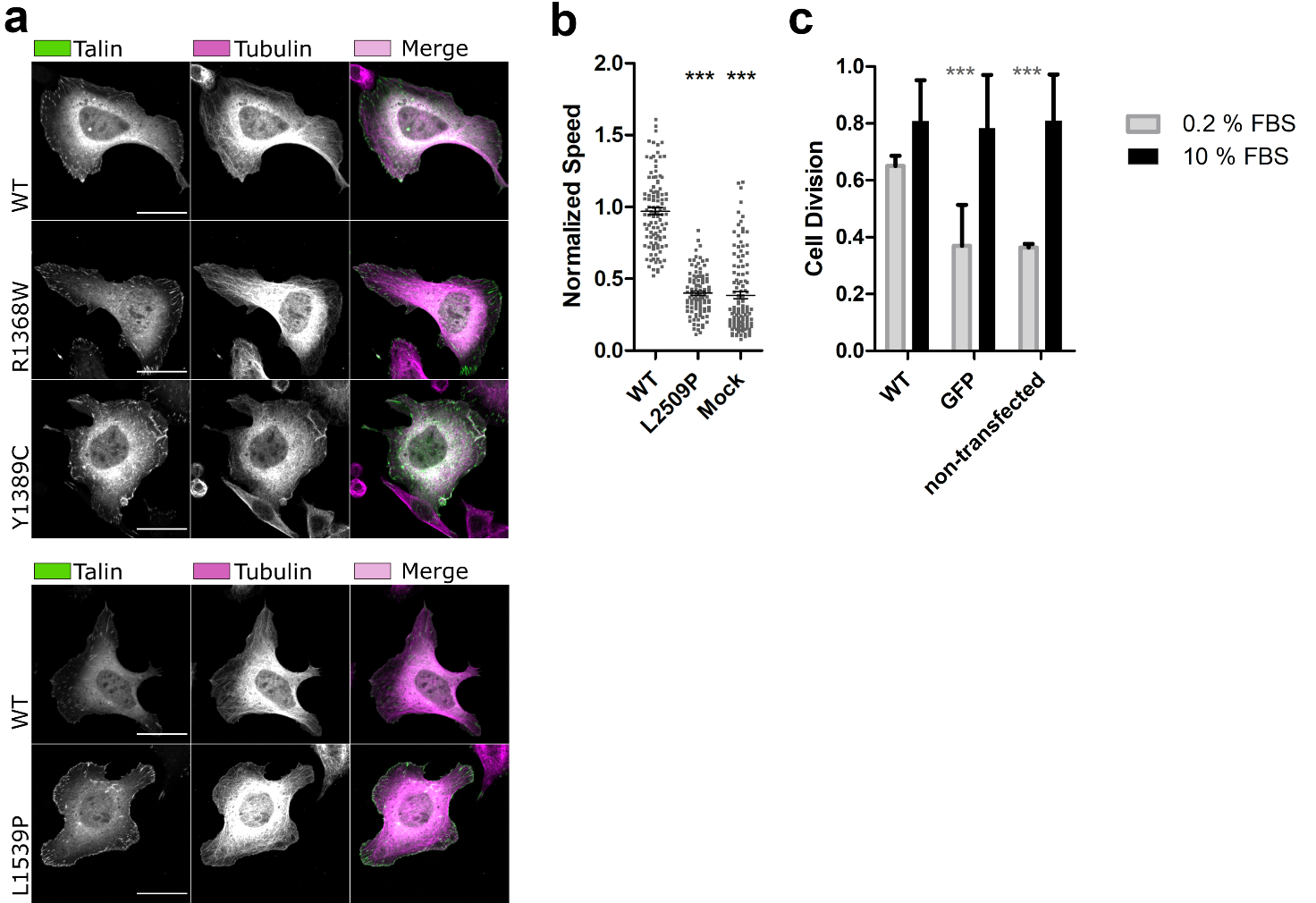

**Figure S5:** (a) SUM projections of z–stacks of cells expressing GFP–tagged talin–1 (WT and/or point mutated) and immunolabeled against Tubulin, ~30 cells per label have been analysed (Fig.5g). (b) Random migration speed (µm/min) determined from time–lapse images of talin expressing cells. Control cells were mock–transfected with GFP–expressing plasmid. The values are normalised to WT and statistical significance measured in comparison to WT. Data are mean +/–SEM. The statistical significance was analysed by one–way ANOVA and Bonferroni test: *P<0.05, **P<0.01, ***P<0.001. (c) Cell division analysis in the presence of 10% FBS and 0.2% FBS defined by the average number of times the cells divide in 12 hours; GFP = transfected mock cells with empty vector; n~ 100 cells from three separate experiment. The statistical analysis was calculated by t–test, non–parametric test of Mann–Whitney: *P<0.05, **P<0.01, ***P<0.001 in comparison to WT for each condition.

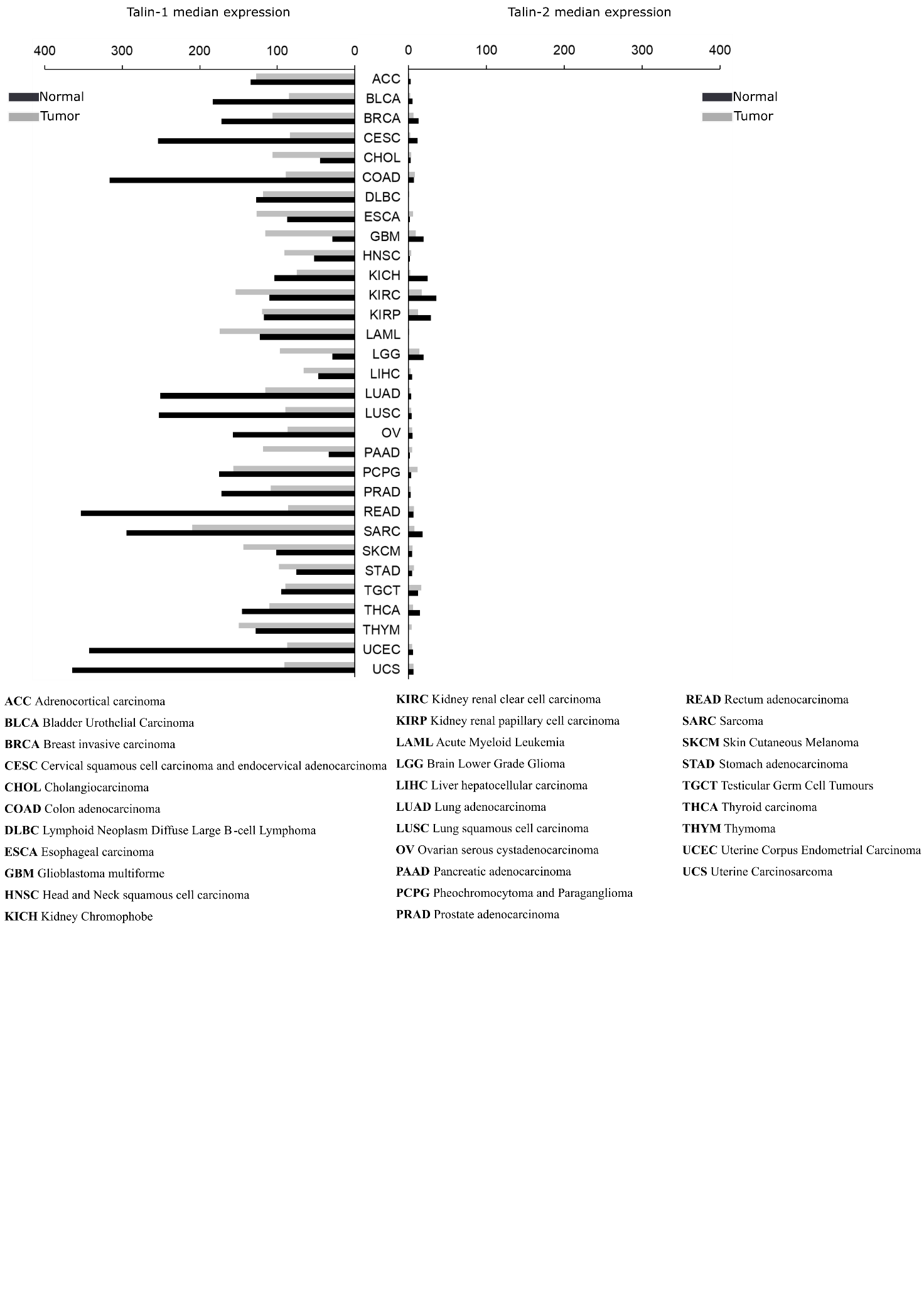

**Figure S6: Talin–1 and talin–2 gene expression profiles across tumour and paired normal samples.** Expression levels taken from GEPIA database, at the time of analysis GEPIA contained 9662 tumour and 5540 normal samples across 33 cancer types.

**Supplementary Materials and Methods**

**Screening TLN1 mutations using bioinformatics**

In order to classify all mutations into different classes and groups, we used structural information from the protein data bank (PDB): 3IVF ^1^, 4F7G ^2^, 1SJ7 ^3^, 2L7A ^4^, 2X0C ^5^, 2KBB ^6^, 3DYJ ^7^ and 2QDQ ^8^. We examined each mutations position on the talin domains and the location on the structure (surface or buried). This was done using an algorithm developed by our group ^9^ and using PyMOL software to visually observe the position of the mutation. The mutations were classified using PON–P2 tool into pathogenic, neutral and unknown to obtain the probability for pathogenicity. PON–P2 is freely available at http://structure.bmc.lu.se/PON–P2/ ^10^. The BLOSUM 62 matrix ^11^ was used to evaluate the severity of the mutations. The degree of evolutionary conservation of the amino acid in the talin–1 sequence was investigated using ConSurf (https://consurf.tau.ac.il/ ^12^). The amino acid substitution matrices CBSM60 was used to analyse the sequence–structure relationship of the protein ^13^. We also investigated whether the mutation was located in known ligand–binding sites and reported the recurrence and the substitution of the amino acid from hydrophobicity to hydrophilicity and vice versa. The mutations were classified into six group based on their location: surface, buried, on a loop, on a linker but buried, between the helices buried, and on the dimerisation domain. These were given the position code of 0.4, 1, 0.2, 1, 0 and 0.8 respectively. For the change in polarity, we gave a score

of one when the amino acid was mutated from hydrophilic residue to hydrophobic residue or from

hydrophobic residue to hydrophilic residue. Considering all these variables, we estimated the scoring coefficient according to the importance feature of each factor using formula n_1_X1+n_2_X2+…+n_n_Xn (n is the scoring coefficient and X is the variable). We tried multiple scoring factor combinations to obtain mutations appearing most frequently with a higher score. Finally, we generated a table from which we selected ten mutants predicted to represent the most drastic mutations based on “total score”. The scoring coefficient to calculate the “total score” of one iteration in Table 1 is as follows: 2*(location within the subdomain code) + 0*(ligand binding code) + 4*(ConSurf code) + 0*(BLOSUM 62) + 2*(PON–P2) 1* (CBSM60) + 0.05*(polarity change from hydrophilic to hydrophobic) + 0.4*(polarity change from hydrophobic to hydrophilic). Altogether, nine iterations were done.

**Prediction of the deleterious effect of mutation**

We normalised the score for each investigated amino–acid substitution between zero and one, with one being the most deleterious. Considering all these factors, we calculated a final score using equation n_1_X1+n_2_X2+…+n_n_Xn by giving different indexes (scoring coefficient, n) a range of zero to five, where a higher value indicated a greater effect of the variant. Using the variables (X), “location within the subdomain”, “ligand binding”, “ConSurf”, “BLOSUM62”, “PON–P2”, “CBSM60” and “polarity change” we tested different relative weightings (for example 2, 0, 4, 0, 2, 1, 0.05, 0.4) and ran nine iterations. Each time, we pooled the top 10 mutations which had the highest score. Based on these criteria, 78 mutants received a score above five, and 10 mutations with the highest scores (>7) likely to be detrimental were taken forward for further analysis.

**Antibodies**

**Table S1:** Antibodies used in this study. Antibodies were diluted in 1.5% BSA, 0.1% Triton–X /PBS buffer. Appropriate secondary antibodies from LI–CORE and a LI–CORE imaging system was used.

| **Antibody** | **Manufacturer** | **Method** | **Dilution used** |
| --- | --- | --- | --- |
| anti–vinculin | Merck, clone hVIN, V9131, RRID:AB_477629 | Immunostaining/  western blot | 1:100 / 1:1000 |
| anti–FAK–pY397 | Abcam, ab81298 [EP2160Y], RRID:AB_1640500 | Immunostaining/  western blot | 1:100 / 1:1000 |
| FAK (clone 77) | BD Biosciences, 610088 | western blot | 1:1000 |
| DLC–1 (H–260) | Santa Cruz Biotechnology, sc–32931 | Immunostaining | 1:100 |
| Integrin β1–chain CD29, clone 9EG7 | BD Pharmingen (Cat:553715) | Immunostaining | 1:200 |
| anti–paxillin | BD Biosciences, 349/Paxillin, 610051, RRID:AB_397463 | Immunostaining/  western blot | 1:100 / 1:1000 |
| GFP antibody | Sicgen AB0020–200 | western blot | 1:1000 |
| Actin | Millipore, MAB 1501R, RRID: AB_2223041 | western blot | 1:2000 |
| Alexa Fluor 568 phalloidin | Life Technologies | Immunostaining | 1:40 |
| Alexa Fluor 568 goat anti–rabbit IgG | Life Technologies A11011 | Immunostaining | 1:200 |
| Alexa Fluor 568 goat anti–mouse IgG | Molecular probes, A11004 | Immunostaining | 1:200 |

**Image analyses**

**Protein expression level and co–localisation quantification from confocal images.** For the quantification of expression level of proteins, the total intensity signal was determined from transfected cells. For the co–localisation quantification of protein intensity, 10–15 adhesion sites per cell were selected based on the EGFP–talin channel using circular selection (0.7 µm) of ImageJ and selection was copied to the red fluorescence channel. Background was assessed from the EGFP channel using circular selections (2.2 µm) from areas devoid of EGFP signal and these areas were again copied to the red fluorescence channel. Protein expression and co–localisation was measured using ImageJ.

**Analyses of adhesion size and number**. To determine the adhesion size and number in transfected cells, we conducted particle analysis using ImageJ particle analyser. First, the threshold range was set to clear out the background noise. Then we selected areas from cell boundaries (roughly one third of the cell membrane) and analysed the adhesions sites based on the signal from the EGFP channel. Cut–off sizes of < 0.1 and > 20 µm^2^ were used in the analyses. Results are shown as ratio of adhesion area and as the number of individual adhesions sites per total selected area per.

**Supplementary** **References**

10. Niroula, A., Urolagin, S. & Vihinen, M. PON-P2: Prediction Method for Fast and Reliable Identification of Harmful Variants. *PloS one* 10, e0117380 (2015).

11. Henikoff, S. & Henikoff, J. G. Amino acid substitution matrices from protein blocks. *Proc. Natl. Acad. Sci. U. S. A.* 89, 10915-10919 (1992).

12. Ashkenazy, H. *et al*. ConSurf 2016: an improved methodology to estimate and visualize evolutionary conservation in macromolecules. *Nucleic Acids Res.* 44, 344 (2016).

13. Liu, X. & Zheng, W. M. An amino acid substitution matrix for protein conformation identification. *J. Bioinform Comput. Biol.* 4, 769-782 (2006).
